# Chromatin remodeling enzyme Snf2h is essential for retinal cell proliferation and photoreceptor maintenance

**DOI:** 10.1101/2023.02.13.528323

**Authors:** Andrea Kuzelova, Naoko Dupacova, Barbora Antosova, Sweetu Susan Sunny, Zbynek Kozmik, Jan Paces, Arthur I. Skoultchi, Tomas Stopka, Zbynek Kozmik

## Abstract

Chromatin remodeling complexes are required for many distinct nuclear processes such as transcription, DNA replication and DNA repair. However, how these complexes contribute to the development of complex tissues within an organism is poorly characterized. Imitation switch (ISWI) proteins are among the most evolutionarily conserved ATP-dependent chromatin remodeling factors and are represented by yeast Isw1/Isw2, and their vertebrate counterparts Snf2h (Smarca5) and Snf2l (Smarca1). In this study, we focused on the role of the *Snf2h* gene during development of the mammalian retina. We show that *Snf2h* is expressed in both retinal progenitors and post-mitotic retinal cells. Using *Snf2h* conditional knockout mice (*Snf2h* cKO), we found that when *Snf2h* is deleted the laminar structure of the adult retina is not retained, the overall thickness of the retina is significantly reduced compared with controls, and the outer nuclear layer (ONL) is completely missing. Depletion of Snf2h did not influence the ability of retinal progenitors to generate all of the differentiated retinal cell types. Instead, Snf2h function is critical for proliferation of retinal progenitor cells. Cells lacking Snf2h have a defective S-phase, leading to the entire cell division process impairments. Although, all retinal cell types appear to be specified in the absence of Snf2h function, cell cycle defects and concomitantly increased apoptosis in *Snf2h* cKO result in abnormal retina lamination, complete destruction of the photoreceptor layer and, consequently, in a physiologically non-functional retina.

## Introduction

In eukaryotes, the chromosomal DNA is compacted in a highly organized nucleoprotein structure called chromatin that presents a barrier to most cellular processes [1]. The nucleosome is the basic structural unit of chromatin and is composed of four histone cores (H2A, H2B, H3 and H4) around which 147 bp of DNA are wrapped [1]. Various ATP-dependent chromatin remodeling complexes can reposition nucleosomes through the energy released by ATP hydrolysis [2,3]. Among the chromatin remodeling ATPases, the imitation switch (ISWI) family is highly conserved during evolution [4–7]. Mammals have two ISWI homologues – Snf2h (sucrose nonfermenting 2 homologue; also known as Smarca 5) and Snf2l (sucrose nonfermenting 2 like; also known as Smarca 1). Both proteins are present in complexes with a diverse array of noncatalytic subunits and are therefore able to promote many biological functions, including DNA replication, DNA repair, transcriptional repression or activation, and maintenance of chromosome structure [8–14].

Snf2h is known to function as the catalytic ATPase in at least five distinct complexes in mammalian cells, including WICH, CHRAC, ACF, RSF and NoRC (reviewed in [3,15]). The presence of Snf2h within these complexes and the interactions of these complexes with other partners results in targeting of Snf2h for particular biological functions in chromatin. For example, Snf2h is recruited to double-strand breaks by sirtuin protein SIRT6 to promote DNA break repair [10]. Lack of SIRT6 and Snf2h impairs chromatin remodeling, increasing sensitivity to genotoxic damage and recruitment of downstream factors such as 53BP1 [10]. Snf2h also plays a major role in organizing arrays of nucleosomes adjacent to the binding sites for the architectural chromatin factor CTCF and acts to promote CTCF binding on DNA [8]. Several studies have focused on the *in vivo* function of Snf2h ATPases and ISWI-containing complexes [6,7,16–23]. However, the simultaneous presence of Snf2h in distinct multicomponent complexes dedicated to unrelated nuclear functions makes it harder to interpret the molecular mechanisms underlying the observed pathological states in the context of a tissue or organism [17,22,24].

There are seven major cell types in mammalian retina that are generated from a common pool of multipotent retinal progenitor cells (RPCs) [25–27]. Each cell type is born in a defined order [26,28–30] and plays a specific role in visual perception [31,32]. Ganglion cells (GCs) arise first, around embryonic day 11 (E11) in mice, and are the only retinal type whose axons project to the brain [31,33,34]. Simultaneously, propagation of other early-born retinal neurons occurs. Horizontal cells (HCs), amacrine cells (ACs) and cone photoreceptors are produced with the highest peak at E14 [35–38]. Thereafter, later-born cells begin to form, namely, bipolar cells (BCs), Müller glia cells (MGCs) and rod photoreceptors, whose generation increases shortly after birth [26,36,38,39]. The process of retinal differentiation is finished at postnatal day 14 (P14) when the eyes open [40]. Mature retinal neurons and glia are arranged in three main layers – ganglion cell layer (GCL), inner nuclear layer (INL) and outer nuclear layer (ONL) [27]. With respect to the entire eye, the GCL is located closest to the lens and is particularly occupied by GCs [36]. The interneurons (HCs, ACs, BCs), primarily associated with transmission of information throughout the retina, are placed together with MGCs, ensuring retinal homeostasis, in the INL [27,36,41]. The ONL, the outermost retinal region, is formed by photoreceptors [36]. Cones and rods are specialized sensory neurons that absorb photons of light and activate the process of photo-transduction [42]. Postnatally, murine photoreceptors express different types of opsin proteins – rhodopsin gene expression is characteristic of rod photoreceptors, while S-opsin (short-wavelength opsins) and M-opsin (medium-wavelength opsins) expression characterizes cone photoreceptors. Photoreceptors are highly metabolically active and require a stable cellular environment, otherwise the cell morphology or physiology is disrupted [43,44]. A number of studies interrogated the role of transcription factors in photoreceptor development [45–56]. In contrast, the regulatory molecules affecting photoreceptor cell maintenance during embryonic and postnatal stages remain largely unknown. It is well established that in order to generate retinal diversity, while maintaining appropriate cell numbers, a proper balance between cell proliferation and differentiation of progenitor cells is required [57,58]. These events are regulated by various extrinsic and intrinsic cues, from which the transcription factors seem to be the most relevant (reviewed in [58,59]). In addition, recent epigenetic studies show that the chromatin-modifying or remodeling mechanisms are also important for retina development and maintenance [60–65]. Here we studied the role of the Snf2h chromatin remodeler during mouse retinal development and differentiation. Since *Snf2h*-deficient mice die at the periimplantation stage due to the growth arrest of trophoectoderm and inner cell mass [22], we performed conditional gene targeting using the floxed allele of *Snf2h* [17]and mRx-Cre active in RPCs from E9.0 onwards [66].

## Materials and methods

### Mouse lines

For the retina-specific inactivation of *Snf2h*, mRx-Cre [66] and Snf2h^fl/fl^ [17] mice were used.

### Tissue collection and histology

Mouse embryos were harvested from timed pregnant females. The morning of vaginal plug was determined as embryonic day 0.5 (E0.5). Embryos and eyes of postnatal mice were fixed in 4% formaldehyde in PBS (w/v) overnight at 4 °C. The next day samples were washed with cold PBS and incubated in 70% ethanol. Subsequently, samples were dehydrated, embedded in paraffin blocks and sectioned. Horizontal sections of 5 μm were prepared, stored at 4°C and used for up to one month.

### Immunohistochemistry

Paraffin sections were deparaffinized and rehydrated. The epitope retrieval was performed for 20 minutes in citrate buffer (10mM, pH 6.0) at 98 °C in a pressure cooker. For immunofluorescent analysis, sections were washed three times in PBT, blocked with 10% BSA in PBT (w/v) for one hour and incubated overnight with primary antibody at 4 °C (diluted in 1% BSA/PBT). Sections were washed three times in PBT, incubated for 45 minutes at room temperature with secondary antibody, washed three times with PBT and covered with DAPI / PBS (1 μg / ml) for 10 minutes. After the wash in PBT, sections were mounted into Mowiol (488; Sigma). For the immunohistological analysis, dewaxed sections were washed three times in PBT, treated with 1.5% H_2_O_2_ in 10% methanol in PBS for 25 minutes, again washed with PBT, blocked with 10% BSA / PBT for one hour and incubated with primary antibody overnight at 4°C. The applied primary antibody was detected with biotinylated secondary antibody (Vector Laboratories) and visualized with Vectastain ABC Elite kit and ImmPACT DAB substrate (all Vector Laboratories). The following primary antibodies were used: rabbit anti-Snf2h (Abcam, 1:500), mouse anti-rhodopsin (Chemicon, 1:200), goat anti-S-opsin (Santa Cruz, 1:500), rabbit anti-M-opsin (Santa Cruz, 1:500), rabbit anti-CENP-A (Cell Signaling, 1:500), rabbit anti-Rxrγ (Santa Cruz, 1:3000), rabbit anti-cCas3 (Cell Signaling, 1:500), goat anti-Lhx2 (Santa Cruz, 1:1000), sheep anti-Chx10 (ThermoFisher Scientific, 1:500), rabbit anti-Otx2 (kindly provided by Dr. Vaccarino, 1:500), rabbit anti-Crx (Santa Cruz, 1:100), rat anti-Blimp1 (Santa Cruz, 1:300), rabbit anti-Pax6 (Covance, 1:500), mouse anti-Lim1/2 (DSHB, 1:250), rabbit anti-Oc1/HNF6 (Santa-Cruz, 1:500), mouse anti-Isl1/2 (DSHB, 1:200), sheep anti-Oc2 (R&D Systems, 1:500), mouse anti-Ap2α (DSHB, 1:1000), rabbit anti-Brn3a (kindly provided by Dr. E. Turner, 1:4000), mouse anti-cyclin D1 (Santa Cruz, 1:300), rabbit anti-pH3 (Santa Cruz, 1:800), rabbit anti-Tbr2 (Abcam, 1:500). Standard histological staining of paraffin sections by hematoxylin and eosin (H&E) was also performed. At least three different embryos from at least two different litters were analyzed with each staining.

### Electroretinography

Overnight dark-adapted mice were anesthetized with ketamine (100 mg/ kg)/ xylazine (20 mg/ kg) and eyes were dilated with a 1% solution (Tropicamide). The entire procedure was performed in a dark room with only a dim red light source. A single flash scotopic electroretinogram (ERG) was recorded using a goldring electrode that was attached to the dilated eye, a reference electrode that was placed in the mouth and a ground electrode punctured in the tail base. Stimuli were produced with a light-emitting diode and the light intensity was gradually amplified during the measurement. The final scotopic ERG waveforms were obtained from P18 wild-type and mRx-Cre; Snf2h^fl/fl^ mice with stimulus intensity 3.0 and 30.0 cd*s/m^2^ (stimulus interval ≥ 60 s).

### EdU incorporation

Timed pregnant females were injected intraperitoneally with 13 μg per g body weight of 5-ethynyl-2’-deoxyuridine (Edu; Invitrogen) and sacrificed after 1 hour or 24 hours. Whole embryos were processed identically as described above. The acquired paraffin sections were deparaffinized, rehydrated and washed three times in PBT. Sections were incubated in proteinase K (20 μg/ml; in TE buffer) at 37 °C for 10 minutes and washed with PBT. Sections were treated with 0.3% H_2_O_2_ in methanol for 20 minutes at room temperature and washed three times in PBT and incubated with Click-iT Edu Imaging kit (AlexaFluor azide, Click iT Edu reaction buffer, CuSO4, Click iT Edu buffer additive) for 1 hour. After rinsing, sections were incubated with DAPI / PBS (1 μg / ml) for 10 minutes, washed in PBT and mounted into Mowiol (4-88; Sigma).

### Quantification of marker-positive cells

The quantification analysis of different retinal cell types, including apoptotic cells and cell cycle analysis, were performed by manual counting of marker-positive cells per central retinal section. In case an eye was removed from the individual (postnatal stages), the eye was precisely oriented into the paraffin block and the central region was verified using a magnifier. Only sections that were kept in place of the optic nerve were taken into account. The number of marker-positive cells per whole retinal section (GCs, HCs) or per defined retinal area in the central retinal part (ACs, BCs, MGCs, rods, cones, apoptotic cells, proliferating cells) was counted and normalized to wild-type controls. For a single eye, a minimum of six sections was used; for each genotype, a minimum of four individual retinae was analyzed. Statistical significance was assessed by Student’s t-test.

### Quantitative RT-PCR (qRT-PCR)

Differentially expressed genes related to the p53 pathway in wild-type and *Snf2h* deficient retinal cells were established in P0 eyes. Postnatal retinae were dissected from the eye, separately from retinal pigment epithelium (RPE) and lens. Total RNA was isolated with Trizol Reagent (Life Technologies). Random-primed cDNA was generated from 500 ng total RNA using the SuperScript VILO cDNA Synthesis Kit (Life Technologies). Six different individuals originating from three litters were used for tissue dissection and subsequent analysis, for both wild-type and *Snf2h*-deficent cells. qRT-PCR reactions were run in a LightCycler 480 Instrument (Roche) using a 5 μl reaction mixture of DNA SYBR Green I Master (Roche) according to the standard manufacturer’s protocol. Analysis was performed on replicates of six different individual samples per genotype, run in triplicate. Crossing point (Cp) values were calculated by LightCycler 480 Software (Roche) using the second-derivate maximum algorithm. The average Cp values of all biological and technical replicates were normalized by Cp values of housekeeping genes. Statistical significance of the change in mRNA expression was calculated by a two-tailed Student’s-test in Microsoft Excel. Finally, the change in mRNA expression was presented as the ratio of mRx-Cre; Snf2h^fl/fl^ to wild-type retinae on a log2 scale. Primer sequences are listed in Table S1.

### RNA-seq

RNA-seq datasets using E14.5, E17.5, P0, P3, P7, P10, P14, and P21 retina from Aldiri et al., 2017[67] were used. FPKM values were obtained from Gene Expression Omunibus (GEO) under accession number GSE87064. Heatmaps were generated using python with standard libraries.

## Results

### Conditional deletion of Snf2h disrupts retinal morphology

To determine *Snf2h* expression during mouse retinal development we performed immunohistochemistry staining stages E9.5 to P18. Snf2h was detected in all RPCs of the optic cup at E9.5 and subsequently in all RPCs and differentiated retinal cells during the later embryonic stages (Fig. 1A-D). After birth, strong *Snf2h* expression was detectable throughout the entire retina, independently of the cell type (Fig. 1I-L).

**Fig. 1.**
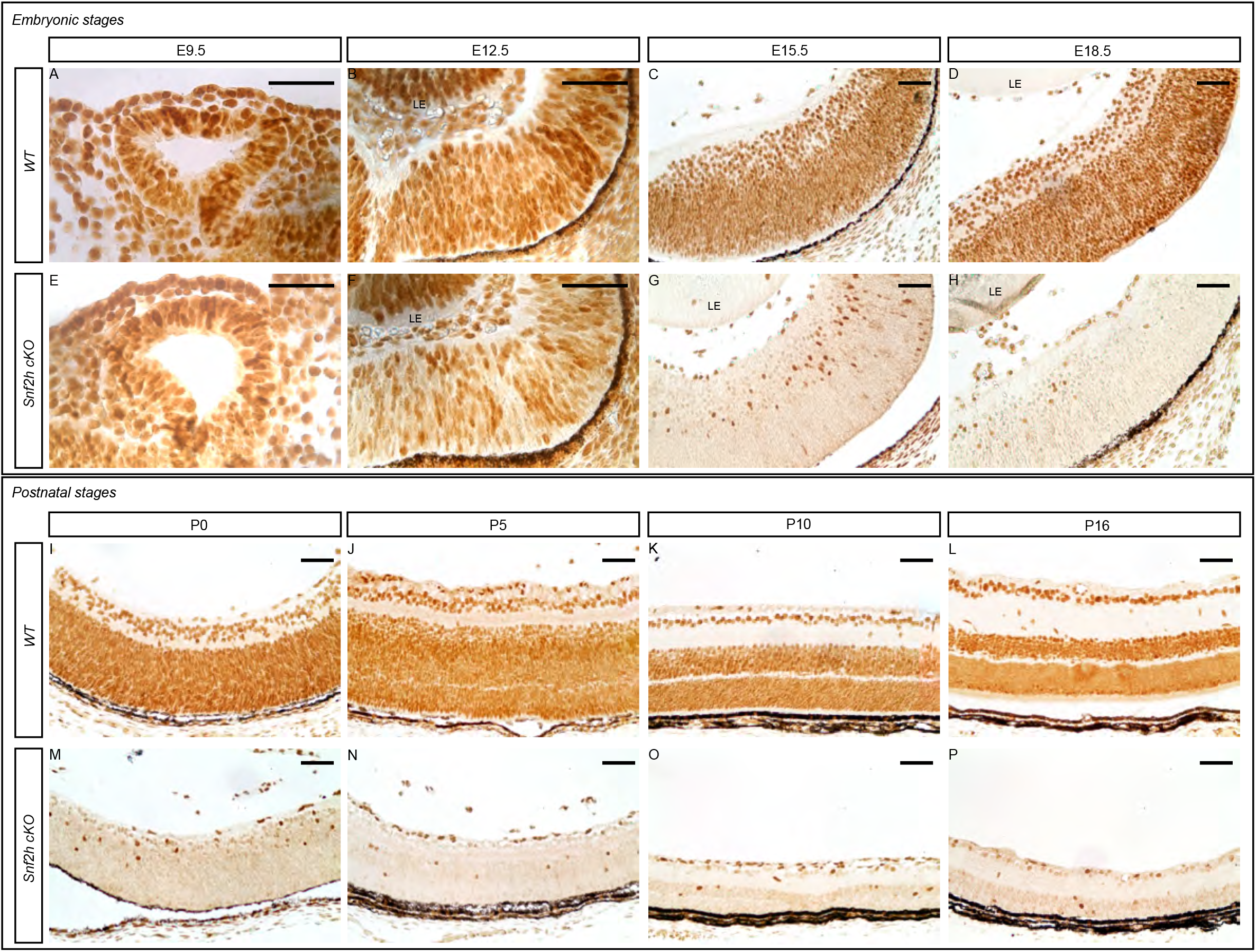
*Snf2h* expression during retina development. Immunohistochemistry staining of wild-type and mRx-Cre; Snf2h^fl/fl^ (*Snf2h* cKO) mice was carried out with anti-Snf2h specific antibody in different embryonic and postnatal stages. *Snf2h* starts to be expressed in early embryonic stages and is maintained in wild-type individuals throughout the embryonic development, and in the adulthood in both retinal progenitors and differentiated retinal cells (A-D, I-L). A decrease of Snf2h levels was first observed in E12.5 conditionally mutant retinae (B, F). During the following embryonic stages, rapid reduction of Snf2h-positive cells in *Snf2h* cKO eye sections occurred (G, H); and only a few Snf2h-positive cells were found in *Snf2h* cKO at postnatal stage (M-P). LE - lens. Scale bars: 50 μm

Snf2h makes at least five ISWI chromatin remodelling complexes (ACF, CHRAC, RSF, WICH, and NoRC)[68–73] and ISWI-complex B-WICH [72,73]. Furthermore, Snf2h interacts with Polycom repressive complex 1 [74], All-1 supercomplex [75], centromere complex [76], Dnmt3b including complexes [77,78], Cohesin complex [79], and HDAC2 [80]. In order to examine gene expression chages of Snf2h and its interacting partners during retinal development, we reanalyzed publicy available RNA-seq data [67]. CORUM database was used to identify components of the complexes [81]. *Snf2h* was remarkably upregulated during P0 and P3 (Fig. S1). Interestingly, majority of ACF, CHRAC, WICH, B-WICH, cohesin, All-1 and Dnmt3b including complex members showed similar expression profiles (Fig. S1-S5).

In order to investigate the function of the *Snf2h* gene during retinal development, we inactivated *Snf2h* conditionally in RPCs and their progeny by crossing mRx-Cre mice with Snf2h^fl/fl^ mice. The mRx-Cre retinal driver is active in RPCs from E9.0 onwards [66], which corresponds with the onset of retinal neurogenesis. We analyzed the extent of Snf2h depletion in the retina of mRx-Cre;Snf2h^fl/fl^ mice (referred to as *Snf2h* cKO) by immunohistochemistry. The first conspicuous reduction of Snf2h-immunoreactivity was observed at E12.5 (Fig. 1F), i.e. at a time when the early-born retinal cell types are already being generated [36]. It is therefore likely that many GCs, HCs, ACs and cones are established in the presence of functional Snf2h protein in the *Snf2h* cKO embryo. During the following embryonic stages examined, the number of retinal cells expressing Snf2h rapidly decreases, and from E15.5 onwards almost all remaining cells lack the Snf2h protein (Fig. 1G, H, M-P). Therefore, some early-born and almost all late-born retinal cells develop in the absence of Snf2h function. To investigate the phenotypic consequences of Snf2h deficiency, we first analyzed the retinal morphology by hematoxylin-eosin (H&E) staining. We did not find any obvious difference between wild-type and *Snf2h* cKO mice in the early embryonic stages (E9.5 – E15.5). The retinal thickness of *Snf2h* cKO was comparable to wild-type control at E16.5 (Fig. 2A, D) and no significant difference was observed until E18.5 (Fig. 2G). After birth, the thickness of the *Snf2h*-deficient retina gradually decreased and at P18 the retina was reduced by 60 % compared with the wild-type control (Fig. 2G). In contrast, *Snf2h* heterozygote mice (genotype mRx-Cre;Snf2h^fl/+^) had normal retinal thickness and morphology even at postnatal week 50 (PW 50, Fig. S6), indicating that a single functional allele of *Snf2h* is sufficient for normal retina development. The reduction of retinal thickness in *Snf2h* cKO was accompanied by selective loss of the outer retinal segment, including ONL and the outer plexiform layer (OPL) (Fig. 2F). It is of note that the retinal lamination was preserved until P16, although the ONL was poorly distinguishable (Fig. 2E). The GCL and INL, along with the inner plexiform layer (IPL), were retained, but thinner compared with wild-type control (Fig. 2C, F).

**Fig. 2.**
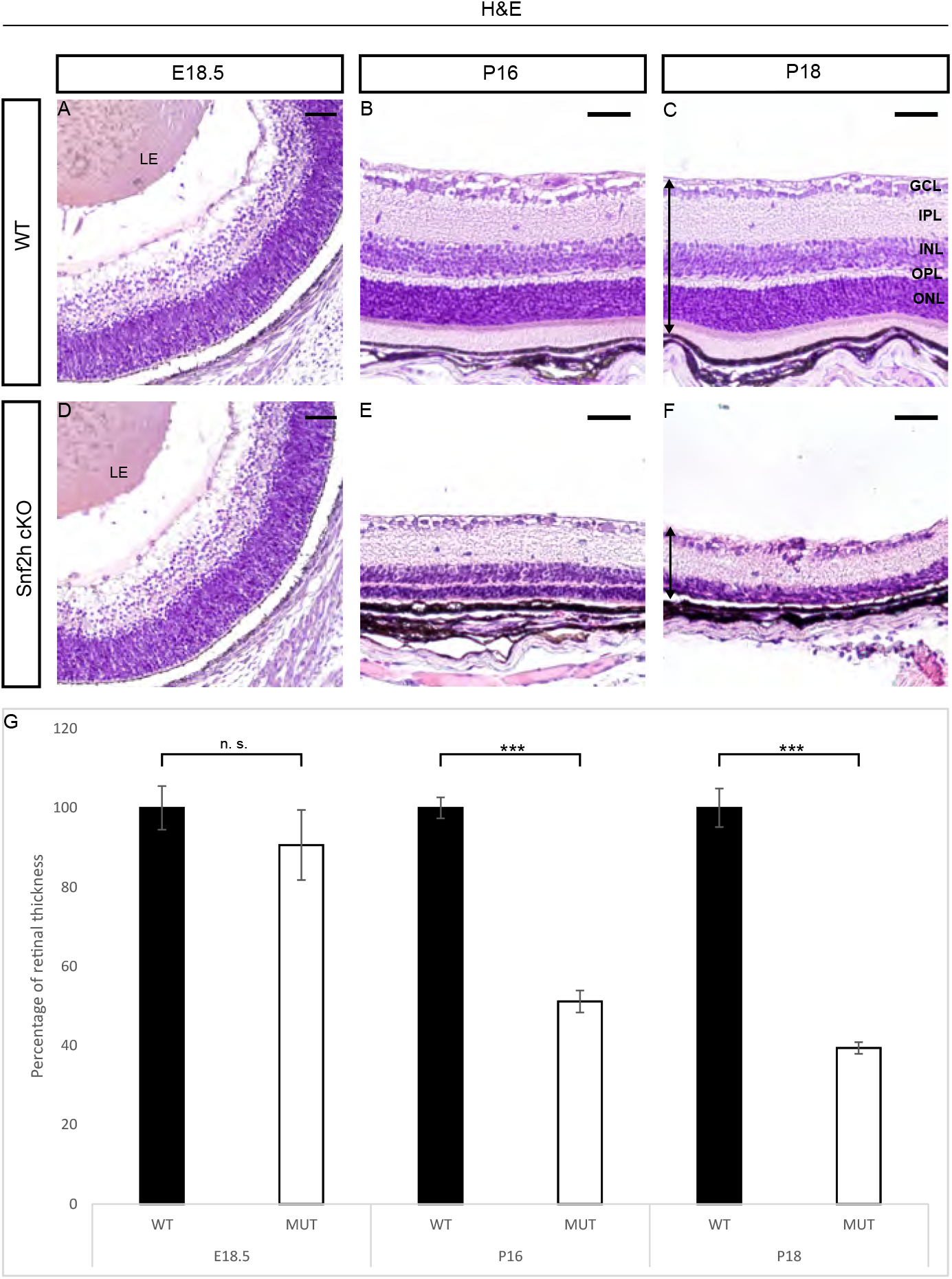
Morphology of *Snf2h*-deficient retina. Hematoxylin and eosin staining of embryonic and postnatal retinal sections of wild-type and *Snf2h* cKO was performed to display the retinal morphology. At the time of fully differentiated retinal cell types (P18), *Snf2h* cKO had dramatically reduced retinal thickness compared with controls (C, F). The outer retinal segment of *Snf2h* cKO mouse was completely missing at P18 (F). *Snf2h*-deficient mice maintained the appropriate retinal structure until P16, although the outer nuclear layer (ONL) was almost indistinguishable (E). Differences in the thickness of the retina were not significant until birth (A, D, G). The time course of the retinal thinning is shown in the graph (G). The arrows show the range from where the retinal thickness was measured (C, F). GCL – ganglion cell layer, INL – inner nuclear layer, ONL – outer nuclear layer, IPL – inner plexiform layer, OPL – outer plexiform layer, LE – lens. Error bars indicate standard derivation, *p*-values are calculated by Student’s t-test (n=3). Scale bars: 50 μm

### Snf2h is required for photoreceptor maintenance and visual perception of mice

Considering that the outermost layer of the retina, which is missing in adult *Snf2h*-deficient mice, is formed by photoreceptors, we analyzed the fate of cones and rods in *Snf2h* cKO mice during embryonic and postnatal stages. To follow photoreceptor specification and maturation, a set of antibodies directed against Otx2, Crx, and Rxrγ was used for photoreceptor mapping during embryonic stages. No significant differences in the immunoreactivity for these photoreceptor markers were observed in *Snf2h* cKO at E18.5 compared with controls (Fig. 3A-F). At P0, we observed an approximately 25% reduction of photoreceptors in *Snf2h* cKO (Fig. 3G-N).

**Fig. 3.**
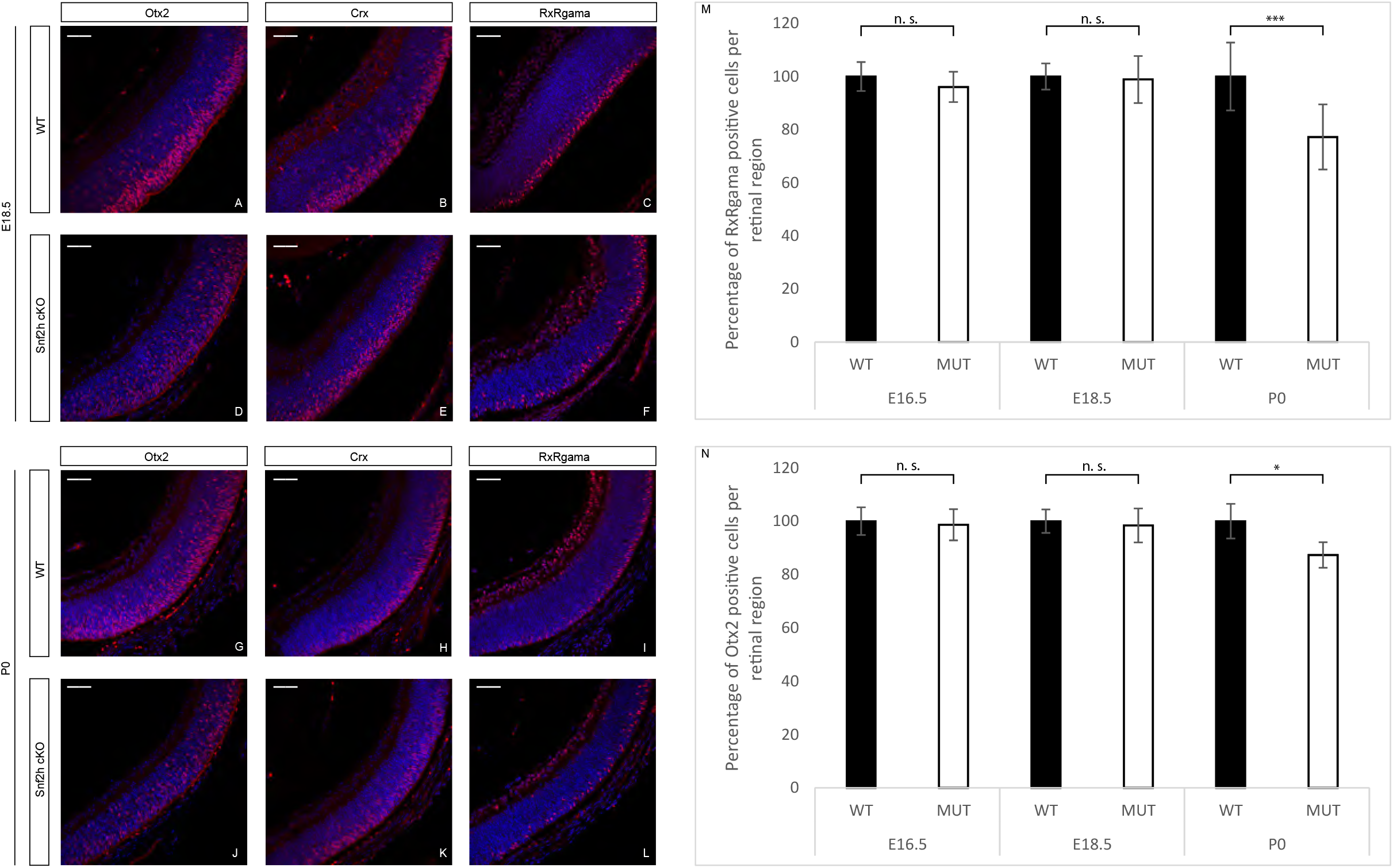
Photoreceptor specification in *Snf2h* cKO. To screen the photoreceptor development, a variety of different photoreceptor-specific markers were used; among them Otx2, Crx, and Rxrγ were chosen for the embryonic immunofluorescent analysis. The immunostained retinal sections showed no significant differences between *Snf2h* cKO mice and controls at E18.5 (A-F). However, there was a rapid decrease of Rxrγ- and Otx2-positive cells in *Snf2h*-deficient mice immediately after birth (G-N). Error bars indicate standard derivation, *p*-values are calculated by Student’s t-test (n=3). Scale bars: 50 μm

At postnatal stages, we used different opsin markers specific for cones or rods, respectively, to follow photoreceptor maturation based on their onset, the level of expression and sub-cellular localization. Rhodopsins are expressed throughout the entire length of the outer retinal segment, whereas M-opsins are preferentially located dorsally, and S-opsins ventrally [55,82]. Since the mouse retina is rod dominated, the prevailing opsin is rhodopsin, which was first detectable in control mice at P5 (Fig. 4A). In *Snf2h* cKO mice, the process of rod photoreceptor maturation appeared to occur normally, although the signal strength of immunostaining was weaker compared with wild-type controls (Fig. 4D). An approximately normal level of rhodopsin expression was detected in *Snf2h*-deficient retina even at P10 (Fig. 4E). Shortly thereafter, the expression of rhodopsin was extinguished, and only sparse rhodopsin-positive cell residues were present at P18 (Fig. 4F). A similar result was obtained by immunostaining for cone photoreceptors. We used two mouse cone-opsin specific markers, Sopsin (short wavelength) and M-opsin (medium wavelength). At P5, the expression of S-opsin was weaker in *Snf2h* cKO mice compared with controls (Fig. 4G, K). At P10, the *Snf2h* deficient retina still retained some S-opsin positive cells, but compared with wild-type their numbers were reduced (Fig. 4I, L). Finally, at P18, the *Snf2h*-deficient cones lacked their characteristic shape and S-opsin positive residues were accumulated just below the INL (Fig. 4J, M). The M-opsin staining at P18 showed a pattern overall similar to that of S-opsin (Fig. 4S). Since the main decline of photoreceptor development took place between P10 and P18, we hypothesized that the key event leading to photoreceptor damage is the eye opening at P14. To test this possibility, we compared *Snf2h* cKO retinas of mice born and raised under normal light conditions (according to parameters in the animal facility) and mice born and raised in the dark. Animals at P13, when the eyes are still closed, and P14, after eye opening, were analyzed. A noticeable difference in the thickness of *Snf2h* cKO retinas between stages P13 and P14 was observed (Fig. S7). Immunodetection for rhodopsin, M- and S-opsin confirmed significant deterioration of morphology of the outer photoreceptor segment between stages P13 (Fig. S8) and P14 (Fig. S9) in *Snf2h* cKO mice. The outer photoreceptor segments were extremely shortened, lost their orientation and spreading within the outermost retinal layer. Our data therefore suggest that an excess of light at the eye opening is not leading factor to photoreceptor damage in *Snf2h* cKO. Instead, the loss of photoreceptors is regulated by intrinsic cues and is gradual.

**Fig. 4.**
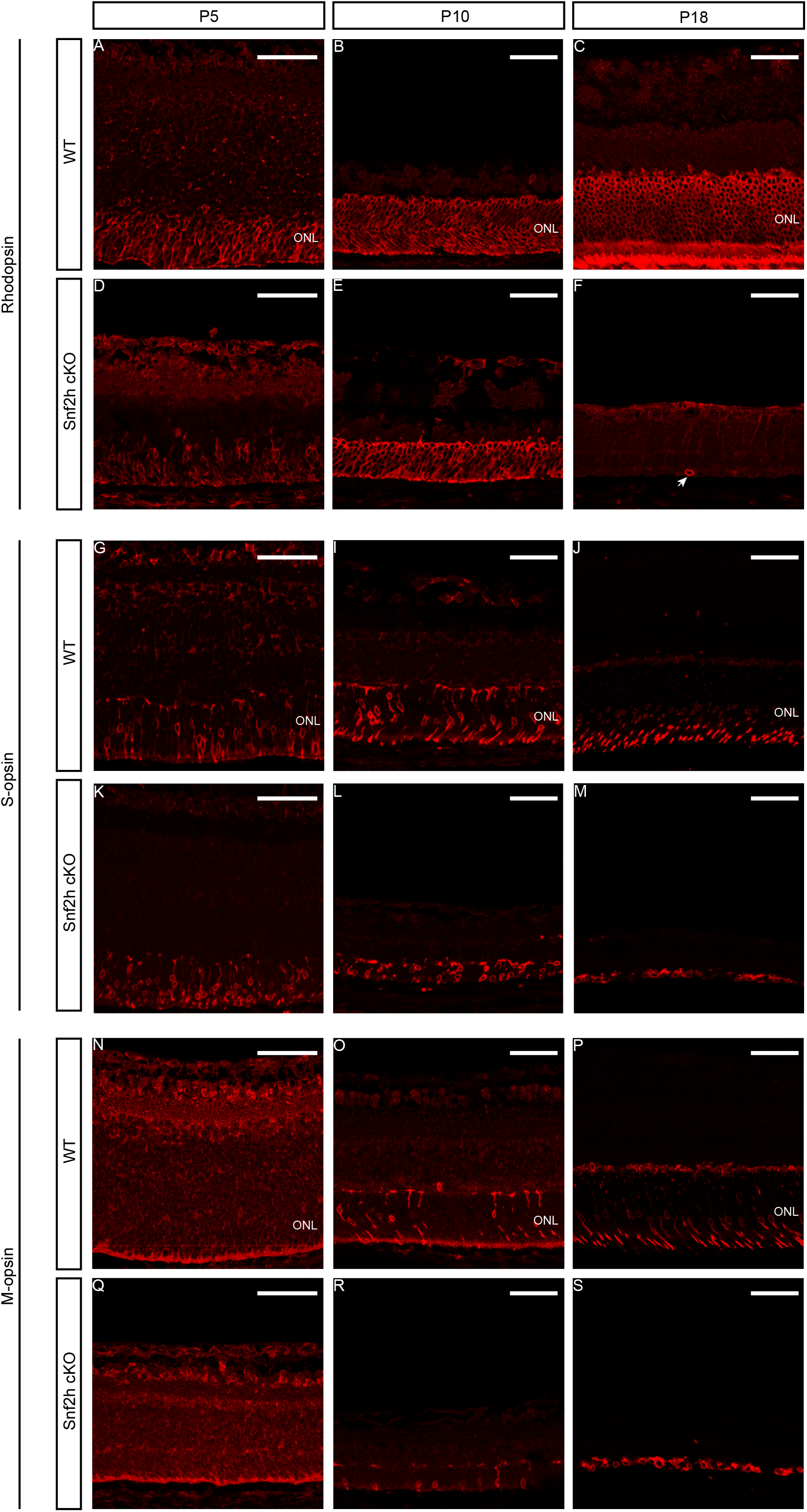
Photoreceptor maintenance in *Snf2h* cKO during postnatal stages. Specific opsin markers were used to determine the photoreceptor state in *Snf2h* cKO after birth. Rhodopsin detects the rod photoreceptors, S-opsin the short-wavelength cone photoreceptors and M-opsin the medium-wavelength cones, respectively. Except for M-opsin that was not expressed until P8, opsin expression was mapped from P5 to P18 (A-S). At P10, all opsins were expressed in *Snf2h-* deficient retinae, although the ONL layer was significantly reduced in thickness compared with controls (B, E; I, L; O, R). At P18, no rhodopsin-positive photoreceptors were present in *Snf2h* cKO, only scattered rhodopsin-positive residues were identified in the entire retinal section (F, indicated by an arrow). At P18, the differences between *Snf2h* cKO and controls in S-opsin (J, M) and M-opsin (P, S) expression were conspicuous, but not as pronounced as the loss of rhodopsin expression. Nevertheless, the shape of cone photoreceptors appeared abnormal. Scale bars: 50 μm

### All other retinal cell types are specified in the absence of Snf2h function

To investigate whether Snf2h function is not restricted to photoreceptors but is also required for generation and/or maintenance of other retinal cell types, we performed immunohistochemistry labelling for specific retinal markers of each cell type. We found that GCs (Brn3a-positive cells, Fig. 5A, F), HCs (Oc2-positive cells, Fig. 5B, G), ACs (Pax6-positive cells, Fig. 5C, H), BCs (Chx10-positive cells, Fig. 5D, I) and MGCs (Lhx2-positive cells, Fig. 5E, J) were present in Snf2h-deficient retina. Combined, our data indicate that *Snf2h* is not necessary for specification of these cell types in the mature mouse retina.

**Fig. 5.**
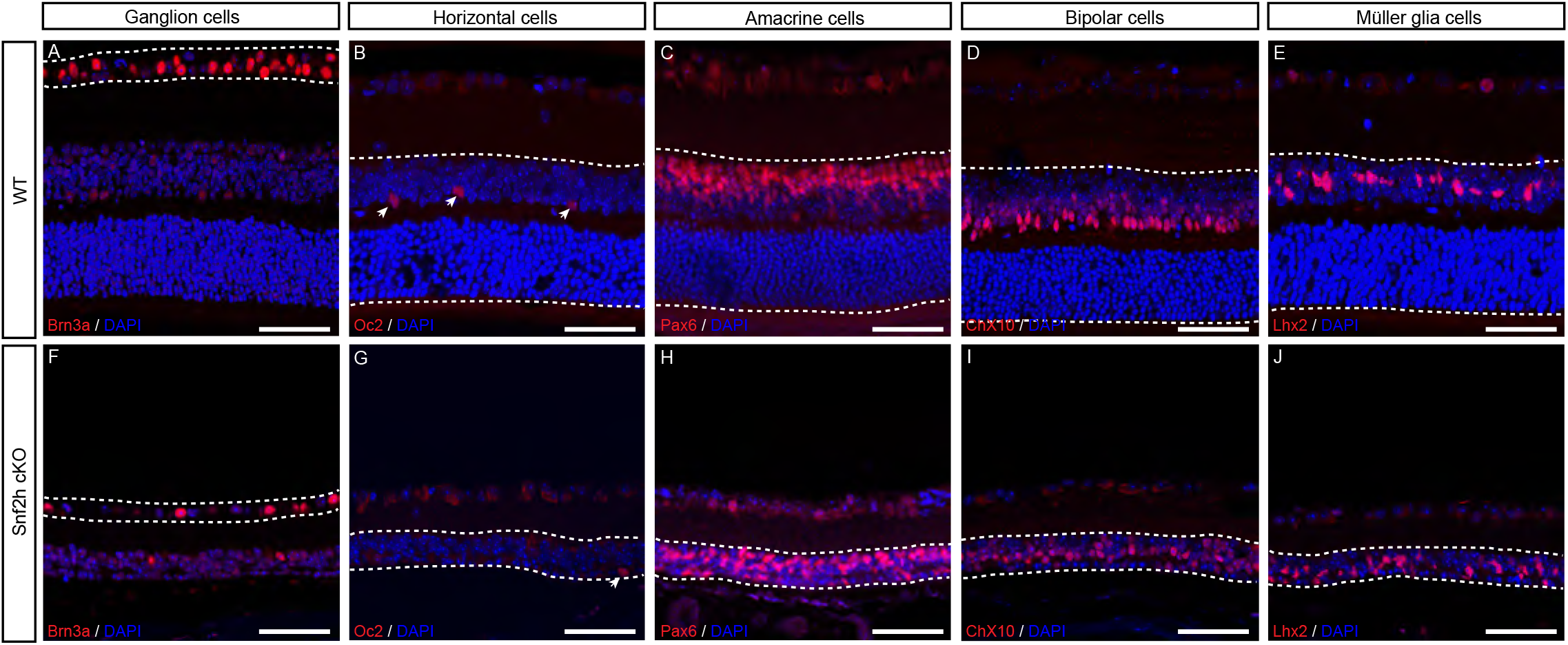
All cell types in GCL and INL are present in *Snf2h*-deficient retina. GCs in the GCL were detected using anti-Brn3a antibodies in P18 retina (A, F). The interneurons in INL of P18 (B, D-E, G, I-J) or P22 (C, H) were identified by the following molecular markers: Oc2 (HCs), Pax6 (ACs), Chx10 (BCs) and Lhx2 (MGCs). Scale bars: 50 μm

### Loss of Snf2h causes cell cycle defects and increased apoptosis

Next, we wanted to determine whether the decreased retinal cell numbers are due to poor RPC expansion or due to initiation of intense apoptosis, or both. To analyze retinal cell proliferation, we examined EdU incorporation during embryonic retina development following a 24hr chase period. The EdU incorporation at P0 revealed dramatic reduction in the number of EdU-positive cells in *Snf2h* cKO - almost no EdU^+^ cells were detectable in contrast to controls, where the proliferation was still high and concentrated in the INL (Fig. 6D, H, I). Such an extreme loss of proliferating cells explains the sharp reduction of the retinal thickness after birth. A major proliferation defect was observed already during embryonic development. The EdU labelling of wild-type and *Snf2h* cKO retinae differed significantly at E16.5 and the difference increased with each consecutive embryonic day (Fig. 6A-H, I). The reduction of EdU-positive cells in *Snf2h*-deficient retina was detectable already at E15.5 following a brief 1h pulse (Fig. 6J-L). Very similar data were obtained using cyclin D1 immunofluorescent labelling (Fig. S10), which does not reflect only the S-phase but rather G1/ S transition. Combined, these results indicated that Snf2h is necessary for proper initiation and progress of DNA replication in retinal cells, which is consistent with previous *in vitro* studies [70,71,83,84]. Next, we used an anti-CENP-A (centromere protein A) antibody recognizing the H3 histone variant that is incorporated into centromeric nucleosomes and is essential for centromere localization and chromosomal segregation. The overall signal was substantially decreased in *Snf2h* cKO mice, in contrast with wild-type animals at E17.5 (Fig. 6M, N). This result suggests that in *Snf2h* cKO retinal cells, the chromosomes are not able to attach to the spindle apparatus, and are therefore not separated. Next, we investigated whether the dramatic decrease of retinal cell numbers is also partly due to increased apoptosis. It was indeed likely that the DNA damage in *Snf2h* cKO leads to chromosomal instability and increased rate of programmed cell death. We first analyzed the apoptosis using an anti-cleaved caspase-3 (cCas3) antibody at E16.5, and in correlation with pronounced proliferation defects, the numbers of apoptotic cells were also increased (Fig. 7A, I). A dramatic increase of cCas3-positive cells was observed in *Snf2h* cKO compared with controls at P0 (Fig. 7D, H, I). It should be noted that both poor replication and increase of apoptosis were not unique to any specific retinal layer, but applied to almost the entire retina. The p53 pathway is often activated upon DNA damage, leading to increased apoptosis. To find out whether the p53 pathway was activated in *Snf2h*-deficient retinae, we analyzed expression of the pathway-related genes by qRT-PCR (Fig. 8). We focused on genes directly controlled by p53, namely *Cdkn1a* (p21), *Cdkn2a* (Arf), *Atm, Atr, Mdm2, Ptf1a* (p48), *Casp3, Casp9, Ccna2* (cyclin A), *Ccnb1* (cyclin B), *Ccne1* (cyclin E) and *Ccng1* (cyclin G). These genes encode proteins involved in various functions: some are necessary for entering the next cell cycle phase, some act as checkpoints, others change their expression levels upon the programmed cell death or directly inhibit p53 function. The qRT-PCR confirmed increased expression of the *Trp53* gene (p53) in *Snf2h* cKO retinae. In addition, the levels of cyclin inhibitor p21 were elevated. This result suggests that the *Snf2h*-depleted retinal cells are arrested in G1 phase of the cell cycle. The altered levels of cyclin E and cyclin B furthermore indicated possible irregularities during the S-phase progression, whereas reduced levels of both Atm and Atr mRNAs in *Snf2h* cKO retinae showed impairments of cell cycle checkpoints. Severe downregulation of cyclin G mRNA levels in *Snf2h*-deficient retinae indicated impaired p53 negative feedback loop, condition that may result in constitutive activation of the p53 pathway. Finally, we compared expression levels of caspase 3 and caspase 9 genes in wild-type and *Snf2h* cKO. Although both genes are expressed during apoptosis, only caspase 3 mRNA levels were upregulated in the mutant retinae.

**Fig. 6.**
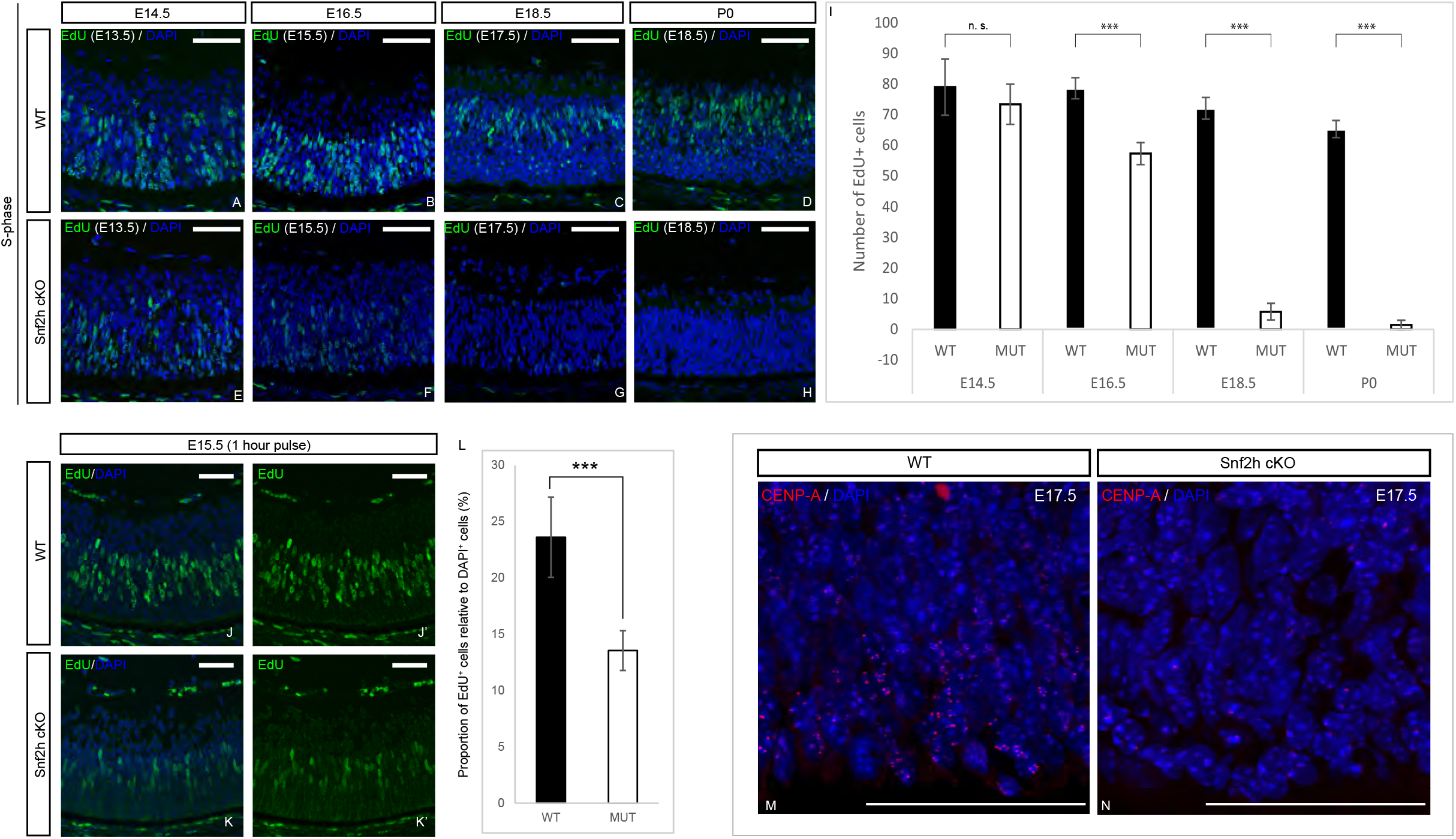
Cell cycle analysis of *Snf2h*-deficient retinal cells. The retinal proliferation was analyzed by EdU incorporation. Pregnant females were intraperitonally injected with EdU 24 hours prior to sacrifice at indicated stages (A-I). The number of EdU^+^ cells in wild-type and *Snf2h* cKO mice was comparable until E14.5 (A, E). The differences first appeared at E16.5 – the number of EdU^+^ cells was decreased by ~ 25 % in *Snf2h*-deficient mice compared with controls (B, F, I). Subsequently, at E18.5 and after birth, almost no EdU^+^ cells were detectable in *Snf2h* cKO, in contrast with the massive proliferation rate in wild-type retinae (C, G; D, H; I). Pregnant females were injected with EdU one hour prior to sacrifice (J-L). The proportion of EdU^+^ cells/DAPI^+^ cells was reduced in *Snf2h*-deficient retina at E15.5 (J-L). Expression of CENP-A (centromere protein A) at E17.5 (M, N). CENP-A is essential for centromere localization and correct chromosomal segregation. The CENP-A antibody staining was widespread in wild-type retinae at E17.5 (M), whereas in *Snf2h*-depleted retinae, only a few CENP-A-positive cells were found (N), indicating impaired chromosomal segregation in *Snf2h* cKO retinal cells. Error bars indicate standard derivation, *p*-values are calculated by Student’s t-test (n=4). Scale bars: 50 μm

**Fig. 7.**
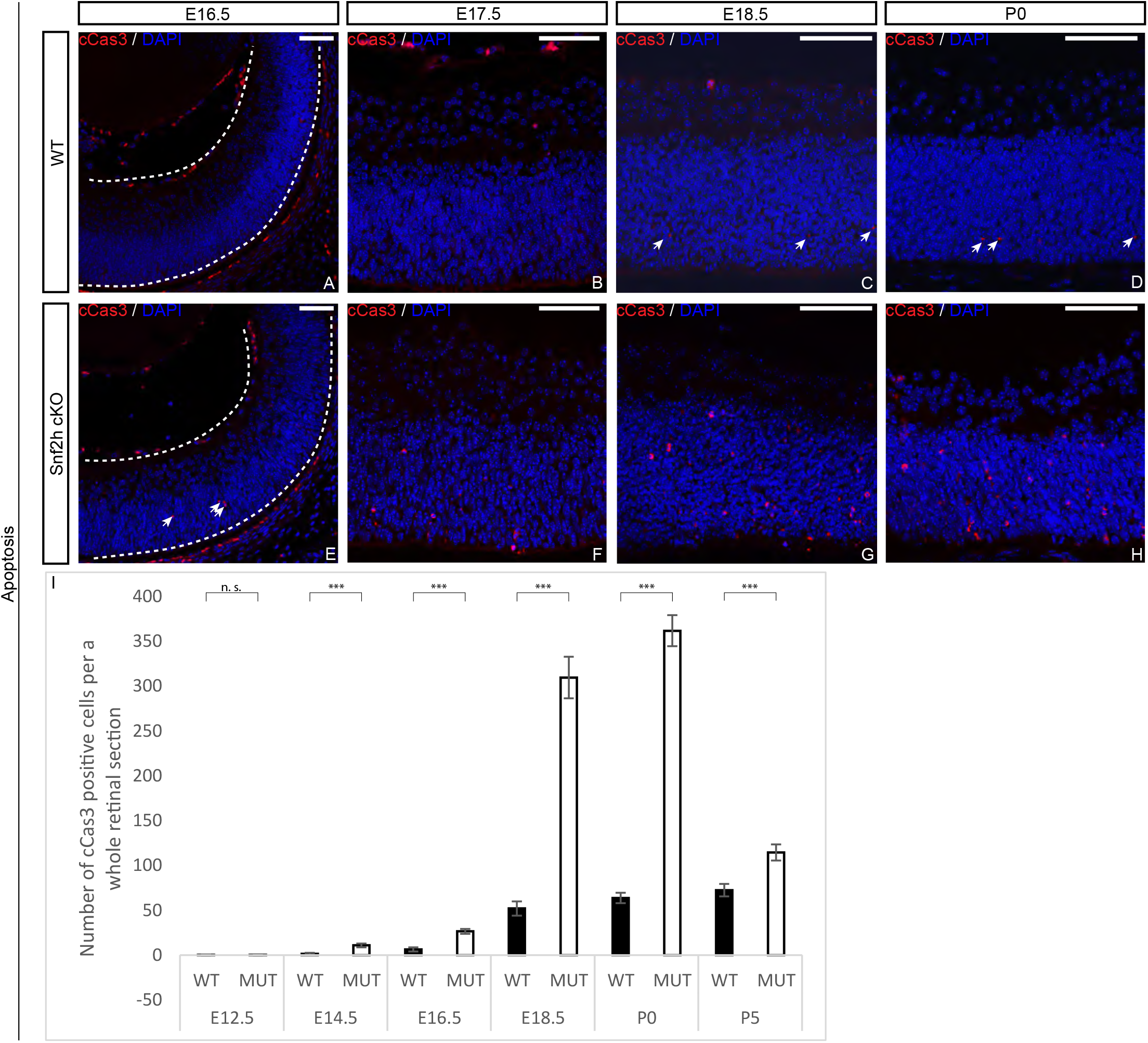
Loss of Snf2h in developing retina is accompanied by massive apoptosis. The number of apoptotic cells in embryonic and postnatal retinae was detected by the anticleaved caspase-3 (cCas3) antibody. The differences between *Snf2h* cKO and controls first appeared at E14.5 (I), and gradually increased during later embryonic stages (A, E; B, F; C, G; I). The peak of apoptosis in *Snf2h*-deficient mice was at P0 (D, H; I). The number of cCas3-positive cells in *Snf2h*-deficient retinae decreased during the postnatal stages; nevertheless, it remained significantly higher at P5 compared with wild-type controls (I). Error bars indicate standard derivation, *p*-values are calculated by Student’s t-test (n=4). Scale bars: 50 μm

**Fig. 8.**
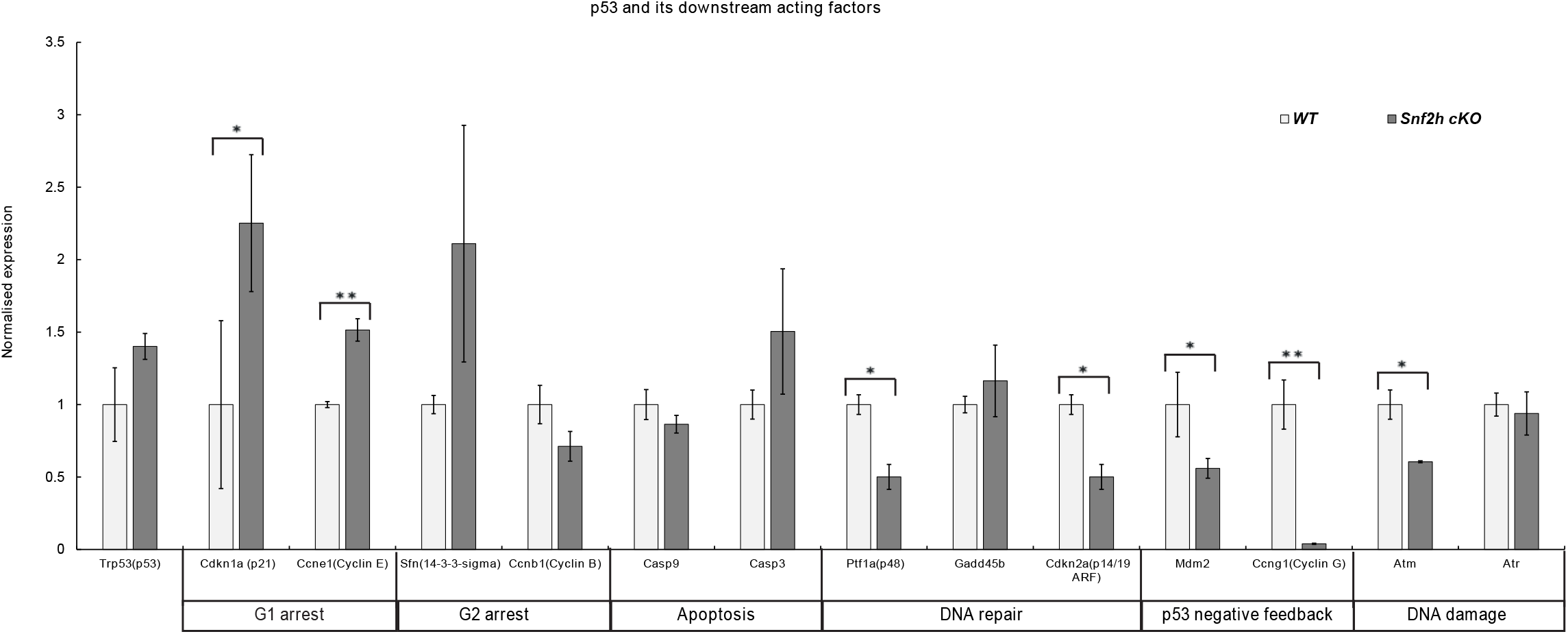
Quantification of mRNA expression of p53 downstream acting factors at P0 assessed by qRT-PCR. Whole retinae from P0 wild-type and *Snf2h* cKO eyes were dissected, subjected to RNA isolation and processed by qRT-PCR. A total of 3 retinas were tested. Error bars indicate standard derivation, *p*-values are calculated by Student’s t-test (n=3).

Taken together our analyses demonstrate cell cycle defects, activation of the p53 pathway mRNA targets and increased apoptosis in *Snf2h* cKO retinae.

## Discussion

The Snf2h protein acts as a chromatin remodeler. Here, through conditional gene targeting and detailed phenotypic analysis, we elucidated the function of the *Snf2h* gene during development, differentiation and maturation of the mouse retina. Considering that Snf2h serves as a catalytic subunit (ATPase) and is incorporated into several distinct ISWI chromatin remodeling complexes (ACF, WICH, CHRAC, RSF, NoRC), the loss of Snf2h likely results in broad impairment of the chromatin dynamics and organization. Snf2h acts as the main executive component of these complexes, whereas the other subunits enhance and direct diverse processes such as DNA replication, DNA repair, recombination and transcription [15,85,86]. Defects observed here in *Snf2h* cKO might be due to the Snf2h function in any of these processes alone or due to combination of defects in several of them. Our data indicated that the *Snf2h*-deficient retina exhibited impaired progenitor cell expansion. Based on EdU incorporation and immunofluorescent staining of proliferation markers, we identified a dominant defect in the S-phase of the cell cycle (process of DNA replication). When *Snf2h* cKO mice were compared with controls, the first remarkable decline in S-phase progression manifested itself at E16.5, i.e. soon after the common peak of production of early-born retinal cells [35–38]. During later embryonic development, the replication activity gradually decreased in *Snf2h* cKO and was minimal at birth. In contrast, in wild-type retinae, proliferation continues approximately up to P8 [30,87,88]. Such a rapid loss of DNA replication was previously observed in the *Snf2h*-depleted Purkinje cells of the mouse cerebellum [17]. Our qRT-PCR results provided further support of aberrant S-phase progression and specifically pointed to the G1/S transition. Increased levels of p21 and cyclin E mRNAs in *Snf2h*-depleted retinal cells indicated cell cycle arrest in G1. It is therefore likely that the extremely rapid decline of DNA replication in embryonic retinal cells is due to the combination of both factors. The importance of Snf2h for DNA replication during the cell cycle was previously shown by several *in vitro* studies [70,71,83,84]. These studies were focused on specific aspects of the S-phase and all established the key role of Snf2h-containing complexes during heterochromatin replication. In addition, the Snf2h function is required in the early S-phase [84], in which the actively transcribed genes within euchromatin are replicated [89]. We propose that the process of cell division is impaired in *Snf2h* cKO retinae. Heterochromatin, whose replication is primarily disrupted, incorporates into the pericentromeric region. This region is characterized as a condensed, transcriptionally repressed chromatin part, necessary for genome stability, chromosome pairing and segregation [90–92]. Perpelescu et al. [93] established a key role of the RSF complex, composed of Rsf1 (remodeling and spacing factor 1) and Snf2h, in maintaining a proper centromere structure by stabilization of the centromere protein A histone variant (CENPA). Our CENP-A analysis indicates that the *Snf2h*-depleted mitotic chromosomes do not contain functional centromeres. It is interesting that Alvarez-Saavedra et al. [17] listed *HJURP* (Holliday Junction Recognition protein) as a gene downregulated in *Snf2h*-deficient cerebellum. HJURP was recently identified as a protein involved in localization of CENP-A into the centromere, and thereby is critical for centromere formation and maintenance [94–97]. If division of the cell nucleus is indeed not properly carried out, then the *Snf2h*-deficient retinal cells likely carry an abnormal number of chromosomes. Hereby, we can conclude that the *Snf2h* gene is required for the proliferation of RPCs. In the absence of Snf2h function, several cell cycle processes occur incorrectly.

Loss of Snf2h in the retina leads to massive apoptosis demonstrated by increased numbers of cleaved caspase 3-positive cells from E16.5 onwards, with a highest peak at P0. Caspase 3 activates the extrinsic apoptotic pathway, whereas caspase 9, whose levels were not significantly changed in the *Snf2h* cKO retinae, activates the intrinsic apoptotic pathway. We speculate that the loss of Snf2h and activation of the p53 pathway influences only the extrinsic apoptotic pathway whose effector caspase is caspase 3.

Given that all retinal cell types are developed from a common pool of retinal progenitor cells, the population of which depletes with proceeding retinogenesis, it is not surprising that the proliferation defects in *Snf2h* cKO cause retina abnormalities in the young adult mice. More remarkable is, however, that the degeneration predominantly affects photoreceptor cells. The reason for this is currently unclear. Cones and rods occupy the outermost layer of the retina and their basic function and morphology is similar despite different birth order during retinogenesis. Cones are generated along with the early-born retinal neurons, whereas rods are classified as later-born cells [36].

The Cre driver used here, the mRx-Cre BAC transgenic line, is an early retina-specific deleter active from E9.0 [66]. In our previous study using this Cre line, we were able to efficiently deplete Pax6 already by E10.5 [98] and Meis1/Meis2 by E14.5 [99]. However, due to the Snf2h protein stability, conspicuous depletion of the Snf2h protein was observed by E12.5 and little to no Snf2h immunoreactivity was present only by E15.5 which is still much earlier compared to the P0 stage by which we observed elimination of the extremely stable polycomb proteins [65]. Considering that the Snf2h depletion in *Snf2h* cKO started at E12.5, many cones were still generated in the presence of Snf2h. This was, however, not the case for late-born rods. Originally, we hypothesized that the birth order of each retinal cell type would play a role in the resulting phenotype of *Snf2h* cKO mice. We assumed that the late-born cell types that develop in the absence of Snf2h function (BCs, MGs, rod photoreceptors) would be much more affected compared with the early-born neurons. Our results indicated, however, that the birth order did not generally correlate with the magnitude of the loss of a particular retinal cell type. GCs, ACs, MGs and BCs were present in adult mice, whereas the photoreceptors had a tendency to disappear completely. Based on marker analysis during embryonic stages (Otx, Crx, Blimp and Rxrγ immunoreactivity) we concluded that the photoreceptors in *Snf2h* cKO were correctly specified and were generated in numbers comparable to wild-type controls. The small loss in photoreceptor numbers occurring before birth was associated with insufficient proliferation and increased apoptosis. Overall, the process of photoreceptor degeneration appeared to be gradual and not directly triggered by external stimuli such as light. Moreover, on closer observation, we found that the photoreceptor collapse could also be caused by damage in synaptogenesis. In case that the photoreceptor cells lose contact with either interneurons or retinal pigment epithelium, a subsequent degenerative process is initiated [100–103]. In case of *Snf2h* cKO, the retinal degeneration occurs early and is fairly rapid. When retinal degeneration occurs, MGCs can activate a gliosis process, which leads to the formation of a glial scar containing a range of auxiliary material [104,105]. This in turn acts as a protection against further neuronal deprivation [106,107]. Such a protective mechanism in *Snf2h*-deficient retina could perhaps contribute to the maintenance of GCL and INL, including the relevant cell types, even in aging mice (at PW 50). Our analysis of *Snf2h*-deficient mice revealed that Snf2h controls expansion of the pool of RPCs by safeguarding the cell cycle progress. In addition, Snf2h appears to be critically required for photoreceptor maintenance in postnatal mouse retina. Since at the molecular level, Snf2h is a catalytic subunit of several distinct multicomponent complexes dedicated to unrelated nuclear processes, it is unlikely that all of the defects observed in *Snf2h* cKO are due to a single Snf2h complex. Further studies aimed at functional characterization of other components of Snf2h containing complexes are needed in order to dissect their specific roles in the retina growth, maturation and maintenance.

## Supplementary Materials

The follwing supporting information can be downloaded at:

## Acknowledgments

We would like to thank Veronika Noskova, Jitka Lachova and Jindra Pohorela for technical assistance. We are grateful to Drs. Vaccarino and Turner for providing reagents and Sarka Takacova for proofreading of the manuscript.

## Author contributions

A.K., N.D. and Z.K.: Conception and project design, acquisition of data, analysis and interpretation of data, drafting and revising the manuscript. B.A., S.S.S., Z.K.Jr., J.P.: Analysis and interpretation of data. A.I.S. and T.S.: provision of Snf2h-floxed mouse strain.

## Institutional Review Board Statement

Housing of mice and in vivo experiments were performed in compliance with the European Communities Council Directive of 24 November 1986 (86/609/EEC) and national and institutional guidelines. Animal care and experimental procedures were approved by the Animal Care Committee of the Institute of Molecular Genetics (no. 71/2014). This work did not include human subjects.

## Data Availability Statement

The data presented in this study are available on request from the corresponding author.

## References

1. Richmond, T.J.; Davey, C.A. The structure of DNA in the nucleosome core. Nature 2003, 423, 145–150, doi:10.1038/nature01595.

2. Narlikar, G.J.; Fan, H.Y.; Kingston, R.E. Cooperation between complexes that regulate chromatin structure and transcription. Cell 2002, 108, 475–487.

3. Jiang, D.; Li, T.; Guo, C.; Tang, T.S.; Liu, H. Small molecule modulators of chromatin remodeling: from neurodevelopment to neurodegeneration. Cell Biosci 2023, 13, 10, doi:10.1186/s13578-023-00953-4.

4. Flaus, A.; Owen-Hughes, T. Mechanisms for ATP-dependent chromatin remodelling: the means to the end. The FEBS journal 2011, 278, 3579–3595, doi:10.1111/j.1742-4658.2011.08281.x.

5. Corona, D.F.; Tamkun, J.W. Multiple roles for ISWI in transcription, chromosome organization and DNA replication. Biochimica et biophysica acta 2004, 1677, 113–119, doi:10.1016/j.bbaexp.2003.09.018.

6. Lazzaro, M.A.; Picketts, D.J. Cloning and characterization of the murine Imitation Switch (ISWI) genes: differential expression patterns suggest distinct developmental roles for Snf2h and Snf2l. Journal of neurochemistry 2001, 77, 1145–1156.

7. Kent, N.A.; Karabetsou, N.; Politis, P.K.; Mellor, J. In vivo chromatin remodeling by yeast ISWI homologs Isw1p and Isw2p. Genes & development 2001, 15, 619–626, doi:10.1101/gad.190301.

8. Wiechens, N.; Singh, V.; Gkikopoulos, T.; Schofield, P.; Rocha, S.; Owen-Hughes, T. The Chromatin Remodelling Enzymes SNF2H and SNF2L Position Nucleosomes adjacent to CTCF and Other Transcription Factors. PLoS genetics 2016, 12, e1005940, doi:10.1371/journal.pgen.1005940.

9. Atsumi, Y.; Minakawa, Y.; Ono, M.; Dobashi, S.; Shinohe, K.; Shinohara, A.; Takeda, S.; Takagi, M.; Takamatsu, N.; Nakagama, H.; et al. ATM and SIRT6/SNF2H Mediate Transient H2AX Stabilization When DSBs Form by Blocking HUWE1 to Allow Efficient gammaH2AX Foci Formation. Cell reports 2015, 13, 2728–2740, doi:10.1016/j.celrep.2015.11.054.

10. Toiber, D.; Erdel, F.; Bouazoune, K.; Silberman, D.M.; Zhong, L.; Mulligan, P.; Sebastian, C.; Cosentino, C.; Martinez-Pastor, B.; Giacosa, S.; et al. SIRT6 recruits SNF2H to DNA break sites, preventing genomic instability through chromatin remodeling. Molecular cell 2013, 51, 454–468, doi:10.1016/j.molcel.2013.06.018.

11. Eberharter, A.; Becker, P.B. ATP-dependent nucleosome remodelling: factors and functions. Journal of cell science 2004, 117, 3707–3711, doi:10.1242/jcs.01175.

12. Kadam, S.; Emerson, B.M. Mechanisms of chromatin assembly and transcription. Current opinion in cell biology 2002, 14, 262–268.

13. Langst, G.; Becker, P.B. Nucleosome mobilization and positioning by ISWI-containing chromatin-remodeling factors. Journal of cell science 2001, 114, 2561–2568.

14. Reyes, A.A.; Marcum, R.D.; He, Y. Structure and Function of Chromatin Remodelers. J Mol Biol 2021, 433, 166929, doi:10.1016/j.jmb.2021.166929.

15. Erdel, F.; Rippe, K. Chromatin remodelling in mammalian cells by ISWI-type complexes--where, when and why? The FEBS journal 2011, 278, 3608–3618, doi:10.1111/j.1742-4658.2011.08282.x.

16. Acemel, R.D.; Tena, J.J.; Irastorza-Azcarate, I.; Marletaz, F.; Gomez-Marin, C.; de la Calle-Mustienes, E.; Bertrand, S.; Diaz, S.G.; Aldea, D.; Aury, J.M.; et al. A single three-dimensional chromatin compartment in amphioxus indicates a stepwise evolution of vertebrate Hox bimodal regulation. Nat Genet 2016, 48, 336–341, doi:10.1038/ng.3497.

17. Alvarez-Saavedra, M.; De Repentigny, Y.; Lagali, P.S.; Raghu Ram, E.V.; Yan, K.; Hashem, E.; Ivanochko, D.; Huh, M.S.; Yang, D.; Mears, A.J.; et al. Snf2h-mediated chromatin organization and histone H1 dynamics govern cerebellar morphogenesis and neural maturation. Nature communications 2014, 5, 4181, doi:10.1038/ncomms5181.

18. Pessina, F.; Lowndes, N.F. The RSF1 histone-remodelling factor facilitates DNA double-strand break repair by recruiting centromeric and Fanconi Anaemia proteins. PLoS biology 2014, 12, e1001856, doi:10.1371/journal.pbio.1001856.

19. Sala, A.; Toto, M.; Pinello, L.; Gabriele, A.; Di Benedetto, V.; Ingrassia, A.M.; Lo Bosco, G.; Di Gesu, V.; Giancarlo, R.; Corona, D.F. Genomewide characterization of chromatin binding and nucleosome spacing activity of the nucleosome remodelling ATPase ISWI. The EMBO journal 2011, 30, 1766–1777, doi:10.1038/emboj.2011.98.

20. Alenghat, T.; Yu, J.; Lazar, M.A. The N-CoR complex enables chromatin remodeler SNF2H to enhance repression by thyroid hormone receptor. The EMBO journal 2006, 25, 3966–3974, doi:10.1038/sj.emboj.7601280.

21. Dirscherl, S.S.; Henry, J.J.; Krebs, J.E. Neural and eye-specific defects associated with loss of the imitation switch (ISWI) chromatin remodeler in Xenopus laevis. Mechanisms of development 2005, 122, 1157–1170, doi:10.1016/j.mod.2005.08.002.

22. Stopka, T.; Skoultchi, A.I. The ISWI ATPase Snf2h is required for early mouse development. Proceedings of the National Academy of Sciences of the United States of America 2003, 100, 14097–14102, doi:10.1073/pnas.2336105100.

23. Deuring, R.; Fanti, L.; Armstrong, J.A.; Sarte, M.; Papoulas, O.; Prestel, M.; Daubresse, G.; Verardo, M.; Moseley, S.L.; Berloco, M.; et al. The ISWI chromatin-remodeling protein is required for gene expression and the maintenance of higher order chromatin structure in vivo. Molecular cell 2000, 5, 355–365.

24. He, S.; Limi, S.; McGreal, R.S.; Xie, Q.; Brennan, L.A.; Kantorow, W.L.; Kokavec, J.; Majumdar, R.; Hou, H., Jr.; Edelmann, W.; et al. Chromatin remodeling enzyme Snf2h regulates embryonic lens differentiation and denucleation. Development (Cambridge, England) 2016, 143, 1937–1947, doi:10.1242/dev.135285.

25. Cepko, C.L. The patterning and onset of opsin expression in vertebrate retinae. Current opinion in neurobiology 1996, 6, 542–546.

26. Turner, D.L.; Cepko, C.L. A common progenitor for neurons and glia persists in rat retina late in development. Nature 1987, 328, 131–136, doi:10.1038/328131a0.

27. Turner, D.L.; Snyder, E.Y.; Cepko, C.L. Lineage-independent determination of cell type in the embryonic mouse retina. Neuron 1990, 4, 833–845.

28. Rapaport, D.H.; Wong, L.L.; Wood, E.D.; Yasumura, D.; LaVail, M.M. Timing and topography of cell genesis in the rat retina. The Journal of comparative neurology 2004, 474, 304–324, doi:10.1002/cne.20134.

29. Young, R.W. Cell differentiation in the retina of the mouse. The Anatomical record 1985, 212, 199–205, doi:10.1002/ar.1092120215.

30. Young, R.W. Cell proliferation during postnatal development of the retina in the mouse. Brain research 1985, 353, 229–239.

31. Masland, R.H. Neuronal diversity in the retina. Current opinion in neurobiology 2001, 11, 431–436.

32. Wassle, H.; Boycott, B.B. Functional architecture of the mammalian retina. Physiological reviews 1991, 71, 447–480, doi:10.1152/physrev.1991.71.2.447.

33. Zagozewski, J.L.; Zhang, Q.; Eisenstat, D.D. Genetic regulation of vertebrate eye development. Clinical genetics 2014, 86, 453–460, doi:10.1111/cge.12493.

34. Hong, Y.K.; Kim, I.J.; Sanes, J.R. Stereotyped axonal arbors of retinal ganglion cell subsets in the mouse superior colliculus. The Journal of comparative neurology 2011, 519, 1691–1711, doi:10.1002/cne.22595.

35. Cherry, T.J.; Trimarchi, J.M.; Stadler, M.B.; Cepko, C.L. Development and diversification of retinal amacrine interneurons at single cell resolution. Proceedings of the National Academy of Sciences of the United States of America 2009, 106, 9495–9500, doi:10.1073/pnas.0903264106.

36. Livesey, F.J.; Cepko, C.L. Vertebrate neural cell-fate determination: lessons from the retina. Nature reviews. Neuroscience 2001, 2, 109–118, doi:10.1038/35053522.

37. Brown, N.L.; Kanekar, S.; Vetter, M.L.; Tucker, P.K.; Gemza, D.L.; Glaser, T. Math5 encodes a murine basic helix-loop-helix transcription factor expressed during early stages of retinal neurogenesis. Development (Cambridge, England) 1998, 125, 4821–4833.

38. Cepko, C.L.; Austin, C.P.; Yang, X.; Alexiades, M.; Ezzeddine, D. Cell fate determination in the vertebrate retina. Proceedings of the National Academy of Sciences of the United States of America 1996, 93, 589–595.

39. Carter-Dawson, L.D.; LaVail, M.M. Rods and cones in the mouse retina. I. Structural analysis using light and electron microscopy. The Journal of comparative neurology 1979, 188, 245–262, doi:10.1002/cne.901880204.

40. Ohsawa, R.; Kageyama, R. Regulation of retinal cell fate specification by multiple transcription factors. Brain research 2008, 1192, 90–98, doi:10.1016/j.brainres.2007.04.014.

41. Bringmann, A.; Iandiev, I.; Pannicke, T.; Wurm, A.; Hollborn, M.; Wiedemann, P.; Osborne, N.N.; Reichenbach, A. Cellular signaling and factors involved in Muller cell gliosis: neuroprotective and detrimental effects. Progress in retinal and eye research 2009, 28, 423–451, doi:10.1016/j.preteyeres.2009.07.001.

42. Fu, Y.; Yau, K.W. Phototransduction in mouse rods and cones. Pflugers Archiv: European journal of physiology 2007, 454, 805–819, doi:10.1007/s00424-006-0194-y.

43. Brzezinski, J.A.; Reh, T.A. Photoreceptor cell fate specification in vertebrates. Development (Cambridge, England) 2015, 142, 3263–3273, doi:10.1242/dev.127043.

44. Du, J.; Rountree, A.; Cleghorn, W.M.; Contreras, L.; Lindsay, K.J.; Sadilek, M.; Gu, H.; Djukovic, D.; Raftery, D.; Satrustegui, J.; et al. Phototransduction Influences Metabolic Flux and Nucleotide Metabolism in Mouse Retina. The Journal of biological chemistry 2016, 291, 4698–4710, doi:10.1074/jbc.M115.698985.

45. Brzezinski, J.A.t.; Kim, E.J.; Johnson, J.E.; Reh, T.A. Ascl1 expression defines a subpopulation of lineage-restricted progenitors in the mammalian retina. Development (Cambridge, England) 2011, 138, 3519–3531, doi:10.1242/dev.064006.

46. Brzezinski, J.A.t.; Lamba, D.A.; Reh, T.A. Blimp1 controls photoreceptor versus bipolar cell fate choice during retinal development. Development (Cambridge, England) 2010, 137, 619–629, doi:10.1242/dev.043968.

47. Katoh, K.; Omori, Y.; Onishi, A.; Sato, S.; Kondo, M.; Furukawa, T. Blimp1 suppresses Chx10 expression in differentiating retinal photoreceptor precursors to ensure proper photoreceptor development. The Journal of neuroscience: the official journal of the Society for Neuroscience 2010, 30, 6515–6526, doi:10.1523/jneurosci.0771-10.2010.

48. Liu, H.; Etter, P.; Hayes, S.; Jones, I.; Nelson, B.; Hartman, B.; Forrest, D.; Reh, T.A. NeuroD1 regulates expression of thyroid hormone receptor 2 and cone opsins in the developing mouse retina. The Journal of neuroscience: the official journal of the Society for Neuroscience 2008, 28, 749–756, doi:10.1523/jneurosci.4832-07.2008.

49. Livne-Bar, I.; Pacal, M.; Cheung, M.C.; Hankin, M.; Trogadis, J.; Chen, D.; Dorval, K.M.; Bremner, R. Chx10 is required to block photoreceptor differentiation but is dispensable for progenitor proliferation in the postnatal retina. Proceedings of the National Academy of Sciences of the United States of America 2006, 103, 4988–4993, doi:10.1073/pnas.0600083103.

50. Nishida, A.; Furukawa, A.; Koike, C.; Tano, Y.; Aizawa, S.; Matsuo, I.; Furukawa, T. Otx2 homeobox gene controls retinal photoreceptor cell fate and pineal gland development. Nature neuroscience 2003, 6, 1255–1263, doi:10.1038/nn1155.

51. Bessant, D.A.; Payne, A.M.; Mitton, K.P.; Wang, Q.L.; Swain, P.K.; Plant, C.; Bird, A.C.; Zack, D.J.; Swaroop, A.; Bhattacharya, S.S. A mutation in NRL is associated with autosomal dominant retinitis pigmentosa. Nature genetics 1999, 21, 355–356, doi:10.1038/7678.

52. Rehemtulla, A.; Warwar, R.; Kumar, R.; Ji, X.; Zack, D.J.; Swaroop, A. The basic motif-leucine zipper transcription factor Nrl can positively regulate rhodopsin gene expression. Proceedings of the National Academy of Sciences of the United States of America 1996, 93, 191–195, doi:10.1073/pnas.93.1.191.

53. Mears, A.J.; Kondo, M.; Swain, P.K.; Takada, Y.; Bush, R.A.; Saunders, T.L.; Sieving, P.A.; Swaroop, A. Nrl is required for rod photoreceptor development. Nature genetics 2001, 29, 447–452, doi:10.1038/ng774.

54. Swaroop, A.; Xu, J.Z.; Pawar, H.; Jackson, A.; Skolnick, C.; Agarwal, N. A conserved retina-specific gene encodes a basic motif/leucine zipper domain. Proceedings of the National Academy of Sciences of the United States of America 1992, 89, 266–270, doi:10.1073/pnas.89.1.266.

55. Swaroop, A.; Kim, D.; Forrest, D. Transcriptional regulation of photoreceptor development and homeostasis in the mammalian retina. Nature reviews. Neuroscience 2010, 11, 563–576, doi:10.1038/nrn2880.

56. Swain, P.K.; Chen, S.; Wang, Q.L.; Affatigato, L.M.; Coats, C.L.; Brady, K.D.; Fishman, G.A.; Jacobson, S.G.; Swaroop, A.; Stone, E.; et al. Mutations in the cone-rod homeobox gene are associated with the cone-rod dystrophy photoreceptor degeneration. Neuron 1997, 19, 1329–1336, doi:10.1016/s0896-6273(00)80423-7.

57. Hardwick, L.J.; Ali, F.R.; Azzarelli, R.; Philpott, A. Cell cycle regulation of proliferation versus differentiation in the central nervous system. Cell and tissue research 2015, 359, 187–200, doi:10.1007/s00441-014-1895-8.

58. Dyer, M.A.; Cepko, C.L. Regulating proliferation during retinal development. Nature reviews. Neuroscience 2001, 2, 333–342, doi:10.1038/35072555.

59. Diacou, R.; Nandigrami, P.; Fiser, A.; Liu, W.; Ashery-Padan, R.; Cvekl, A. Cell fate decisions, transcription factors and signaling during early retinal development. Progress in retinal and eye research 2022, 91, 101093, doi:10.1016/j.preteyeres.2022.101093.

60. Zhang, J.; Taylor, R.J.; La Torre, A.; Wilken, M.S.; Cox, K.E.; Reh, T.A.; Vetter, M.L. Ezh2 maintains retinal progenitor proliferation, transcriptional integrity, and the timing of late differentiation. Developmental biology 2015, 403, 128–138, doi:10.1016/j.ydbio.2015.05.010.

61. Popova, E.Y.; Grigoryev, S.A.; Fan, Y.; Skoultchi, A.I.; Zhang, S.S.; Barnstable, C.J. Developmentally regulated linker histone H1c promotes heterochromatin condensation and mediates structural integrity of rod photoreceptors in mouse retina. The Journal of biological chemistry 2013, 288, 17895–17907, doi:10.1074/jbc.M113.452144.

62. Popova, E.Y.; Barnstable, C.J.; Zhang, S.S. Cell type-specific epigenetic signatures accompany late stages of mouse retina development. Advances in experimental medicine and biology 2014, 801, 3–8, doi:10.1007/978-1-4614-3209-8_1.

63. Merbs, S.L.; Khan, M.A.; Hackler, L., Jr.; Oliver, V.F.; Wan, J.; Qian, J.; Zack, D.J. Cell-specific DNA methylation patterns of retina-specific genes. PloS one 2012, 7, e32602, doi:10.1371/journal.pone.0032602.

64. Lagali, P.S.; Picketts, D.J. Matters of life and death: the role of chromatin remodeling proteins in retinal neuron survival. Journal of ocular biology, diseases, and informatics 2011, 4, 111–120, doi:10.1007/s12177-012-9080-3.

65. Fujimura, N.; Kuzelova, A.; Ebert, A.; Strnad, H.; Lachova, J.; Machon, O.; Busslinger, M.; Kozmik, Z. Polycomb repression complex 2 is required for the maintenance of retinal progenitor cells and balanced retinal differentiation. Developmental biology 2018, 433, 47–60, doi:10.1016/j.ydbio.2017.11.004.

66. Klimova, L.; Lachova, J.; Machon, O.; Sedlacek, R.; Kozmik, Z. Generation of mRx-Cre transgenic mouse line for efficient conditional gene deletion in early retinal progenitors. PloS one 2013, 8, e63029, doi:10.1371/journal.pone.0063029.

67. Aldiri, I.; Xu, B.; Wang, L.; Chen, X.; Hiler, D.; Griffiths, L.; Valentine, M.; Shirinifard, A.; Thiagarajan, S.; Sablauer, A.; et al. The Dynamic Epigenetic Landscape of the Retina During Development, Reprogramming, and Tumorigenesis. Neuron 2017, 94, 550–568 e510, doi:10.1016/j.neuron.2017.04.022.

68. Strohner, R.; Nemeth, A.; Jansa, P.; Hofmann-Rohrer, U.; Santoro, R.; Langst, G.; Grummt, I. NoRC--a novel member of mammalian ISWI-containing chromatin remodeling machines. The EMBO journal 2001, 20, 4892–4900, doi:10.1093/emboj/20.17.4892.

69. LeRoy, G.; Orphanides, G.; Lane, W.S.; Reinberg, D. Requirement of RSF and FACT for transcription of chromatin templates in vitro. Science 1998, 282, 1900–1904, doi:10.1126/science.282.5395.1900.

70. Bozhenok, L.; Wade, P.A.; Varga-Weisz, P. WSTF-ISWI chromatin remodeling complex targets heterochromatic replication foci. The EMBO journal 2002, 21, 2231–2241, doi:10.1093/emboj/21.9.2231.

71. Poot, R.A.; Dellaire, G.; Hulsmann, B.B.; Grimaldi, M.A.; Corona, D.F.; Becker, P.B.; Bickmore, W.A.; Varga-Weisz, P.D. HuCHRAC, a human ISWI chromatin remodelling complex contains hACF1 and two novel histone-fold proteins. The EMBO journal 2000, 19, 3377–3387, doi:10.1093/emboj/19.13.3377.

72. Cavellan, E.; Asp, P.; Percipalle, P.; Farrants, A.K. The WSTF-SNF2h chromatin remodeling complex interacts with several nuclear proteins in transcription. The Journal of biological chemistry 2006, 281, 16264–16271, doi:10.1074/jbc.M600233200.

73. Sadeghifar, F.; Bohm, S.; Vintermist, A.; Ostlund Farrants, A.K. The B-WICH chromatin-remodelling complex regulates RNA polymerase III transcription by promoting Max-dependent c-Myc binding. Nucleic acids research 2015, 43, 4477–4490, doi:10.1093/nar/gkv312.

74. Levine, S.S.; Weiss, A.; Erdjument-Bromage, H.; Shao, Z.; Tempst, P.; Kingston, R.E. The core of the polycomb repressive complex is compositionally and functionally conserved in flies and humans. Mol Cell Biol 2002, 22, 6070–6078, doi:10.1128/MCB.22.17.6070-6078.2002.

75. Nakamura, T.; Mori, T.; Tada, S.; Krajewski, W.; Rozovskaia, T.; Wassell, R.; Dubois, G.; Mazo, A.; Croce, C.M.; Canaani, E. ALL-1 is a histone methyltransferase that assembles a supercomplex of proteins involved in transcriptional regulation. Molecular cell 2002, 10, 1119–1128, doi:10.1016/s1097-2765(02)00740-2.

76. Obuse, C.; Yang, H.; Nozaki, N.; Goto, S.; Okazaki, T.; Yoda, K. Proteomics analysis of the centromere complex from HeLa interphase cells: UV-damaged DNA binding protein 1 (DDB-1) is a component of the CEN-complex, while BMI-1 is transiently co-localized with the centromeric region in interphase. Genes Cells 2004, 9, 105–120, doi:10.1111/j.1365-2443.2004.00705.x.

77. Geiman, T.M.; Sankpal, U.T.; Robertson, A.K.; Chen, Y.; Mazumdar, M.; Heale, J.T.; Schmiesing, J.A.; Kim, W.; Yokomori, K.; Zhao, Y.; et al. Isolation and characterization of a novel DNA methyltransferase complex linking DNMT3B with components of the mitotic chromosome condensation machinery. Nucleic acids research 2004, 32, 2716–2729, doi:10.1093/nar/gkh589.

78. Zhou, Y.; Grummt, I. The PHD finger/bromodomain of NoRC interacts with acetylated histone H4K16 and is sufficient for rDNA silencing. Current biology: CB 2005, 15, 1434–1438, doi:10.1016/j.cub.2005.06.057.

79. Hakimi, M.A.; Bochar, D.A.; Schmiesing, J.A.; Dong, Y.; Barak, O.G.; Speicher, D.W.; Yokomori, K.; Shiekhattar, R. A chromatin remodelling complex that loads cohesin onto human chromosomes. Nature 2002, 418, 994–998, doi:10.1038/nature01024.

80. Zhou, J.; Chau, C.M.; Deng, Z.; Shiekhattar, R.; Spindler, M.P.; Schepers, A.; Lieberman, P.M. Cell cycle regulation of chromatin at an origin of DNA replication. The EMBO journal 2005, 24, 1406–1417, doi:10.1038/sj.emboj.7600609.

81. Tsitsiridis, G.; Steinkamp, R.; Giurgiu, M.; Brauner, B.; Fobo, G.; Frishman, G.; Montrone, C.; Ruepp, A. CORUM: the comprehensive resource of mammalian protein complexes-2022. Nucleic acids research 2023, 51, D539–D545, doi:10.1093/nar/gkac1015.

82. Applebury, M.L.; Antoch, M.P.; Baxter, L.C.; Chun, L.L.; Falk, J.D.; Farhangfar, F.; Kage, K.; Krzystolik, M.G.; Lyass, L.A.; Robbins, J.T. The murine cone photoreceptor: a single cone type expresses both S and M opsins with retinal spatial patterning. Neuron 2000, 27, 513–523.

83. Poot, R.A.; Bozhenok, L.; van den Berg, D.L.; Steffensen, S.; Ferreira, F.; Grimaldi, M.; Gilbert, N.; Ferreira, J.; Varga-Weisz, P.D. The Williams syndrome transcription factor interacts with PCNA to target chromatin remodelling by ISWI to replication foci. Nature cell biology 2004, 6, 1236–1244, doi:10.1038/ncb1196.

84. Collins, N.; Poot, R.A.; Kukimoto, I.; Garcia-Jimenez, C.; Dellaire, G.; Varga-Weisz, P.D. An ACF1-ISWI chromatin-remodeling complex is required for DNA replication through heterochromatin. Nature genetics 2002, 32, 627–632, doi:10.1038/ng1046.

85. Eberharter, A.; Ferrari, S.; Langst, G.; Straub, T.; Imhof, A.; Varga-Weisz, P.; Wilm, M.; Becker, P.B. Acf1, the largest subunit of CHRAC, regulates ISWI-induced nucleosome remodelling. The EMBO journal 2001, 20, 3781–3788, doi:10.1093/emboj/20.14.3781.

86. Ito, T.; Levenstein, M.E.; Fyodorov, D.V.; Kutach, A.K.; Kobayashi, R.; Kadonaga, J.T. ACF consists of two subunits, Acf1 and ISWI, that function cooperatively in the ATP-dependent catalysis of chromatin assembly. Genes & development 1999, 13, 1529–1539.

87. Zhang, J.; Gray, J.; Wu, L.; Leone, G.; Rowan, S.; Cepko, C.L.; Zhu, X.; Craft, C.M.; Dyer, M.A. Rb regulates proliferation and rod photoreceptor development in the mouse retina. Nature genetics 2004, 36, 351–360, doi:10.1038/ng1318.

88. Alexiades, M.R.; Cepko, C. Quantitative analysis of proliferation and cell cycle length during development of the rat retina. Developmental dynamics: an official publication of the American Association of Anatomists 1996, 205, 293–307, doi:10.1002/(sici)1097-0177(199603)205:3<293::aid-aja9>3.0.co;2-d.

89. O’Keefe, R.T.; Henderson, S.C.; Spector, D.L. Dynamic organization of DNA replication in mammalian cell nuclei: spatially and temporally defined replication of chromosome-specific alpha-satellite DNA sequences. The Journal of cell biology 1992, 116, 1095–1110.

90. Bernard, P.; Allshire, R. Centromeres become unstuck without heterochromatin. Trends in cell biology 2002, 12, 419–424.

91. Taddei, A.; Maison, C.; Roche, D.; Almouzni, G. Reversible disruption of pericentric heterochromatin and centromere function by inhibiting deacetylases. Nature cell biology 2001, 3, 114–120, doi:10.1038/35055010.

92. Wallrath, L.L. Unfolding the mysteries of heterochromatin. Current opinion in genetics & development 1998, 8, 147–153.

93. Perpelescu, M.; Nozaki, N.; Obuse, C.; Yang, H.; Yoda, K. Active establishment of centromeric CENP-A chromatin by RSF complex. The Journal of cell biology 2009, 185, 397–407, doi:10.1083/jcb.200903088.

94. Perpelescu, M.; Hori, T.; Toyoda, A.; Misu, S.; Monma, N.; Ikeo, K.; Obuse, C.; Fujiyama, A.; Fukagawa, T. HJURP is involved in the expansion of centromeric chromatin. Molecular biology of the cell 2015, 26, 2742–2754, doi:10.1091/mbc.E15-02-0094.

95. Tachiwana, H.; Muller, S.; Blumer, J.; Klare, K.; Musacchio, A.; Almouzni, G. HJURP involvement in de novo CenH3(CENP-A) and CENP-C recruitment. Cell reports 2015, 11, 22–32, doi:10.1016/j.celrep.2015.03.013.

96. Bassett, E.A.; DeNizio, J.; Barnhart-Dailey, M.C.; Panchenko, T.; Sekulic, N.; Rogers, D.J.; Foltz, D.R.; Black, B.E. HJURP uses distinct CENP-A surfaces to recognize and to stabilize CENP-A/histone H4 for centromere assembly. Developmental cell 2012, 22, 749–762, doi:10.1016/j.devcel.2012.02.001.

97. Foltz, D.R.; Jansen, L.E.; Bailey, A.O.; Yates, J.R., 3rd; Bassett, E.A.; Wood, S.; Black, B.E.; Cleveland, D.W. Centromere-specific assembly of CENP-a nucleosomes is mediated by HJURP. Cell 2009, 137, 472–484, doi:10.1016/j.cell.2009.02.039.

98. Klimova, L.; Kozmik, Z. Stage-dependent requirement of neuroretinal Pax6 for lens and retina development. Development (Cambridge, England) 2014, 141, 1292–1302, doi:10.1242/dev.098822.

99. Dupacova, N.; Antosova, B.; Paces, J.; Kozmik, Z. Meis homeobox genes control progenitor competence in the retina. Proceedings of the National Academy of Sciences of the United States of America 2021, 118, doi:10.1073/pnas.2013136118.

100. Housset, M.; Samuel, A.; Ettaiche, M.; Bemelmans, A.; Beby, F.; Billon, N.; Lamonerie, T. Loss of Otx2 in the adult retina disrupts retinal pigment epithelium function, causing photoreceptor degeneration. The Journal of neuroscience: the official journal of the Society for Neuroscience 2013, 33, 9890–9904, doi:10.1523/jneurosci.1099-13.2013.

101. Sonntag, S.; Dedek, K.; Dorgau, B.; Schultz, K.; Schmidt, K.F.; Cimiotti, K.; Weiler, R.; Lowel, S.; Willecke, K.; Janssen-Bienhold, U. Ablation of retinal horizontal cells from adult mice leads to rod degeneration and remodeling in the outer retina. The Journal of neuroscience: the official journal of the Society for Neuroscience 2012, 32, 10713–10724, doi:10.1523/jneurosci.0442-12.2012.

102. Pacione, L.R.; Szego, M.J.; Ikeda, S.; Nishina, P.M.; McInnes, R.R. Progress toward understanding the genetic and biochemical mechanisms of inherited photoreceptor degenerations. Annual review of neuroscience 2003, 26, 657–700, doi:10.1146/annurev.neuro.26.041002.131416.

103. Blanks, J.C.; Adinolfi, A.M.; Lolley, R.N. Photoreceptor degeneration and synaptogenesis in retinal-degenerative (rd) mice. The Journal of comparative neurology 1974, 156, 95–106, doi:10.1002/cne.901560108.

104. Jones, B.W.; Watt, C.B.; Frederick, J.M.; Baehr, W.; Chen, C.K.; Levine, E.M.; Milam, A.H.; Lavail, M.M.; Marc, R.E. Retinal remodeling triggered by photoreceptor degenerations. The Journal of comparative neurology 2003, 464, 1–16, doi:10.1002/cne.10703.

105. Jones, B.W.; Marc, R.E. Retinal remodeling during retinal degeneration. Experimental eye research 2005, 81, 123–137, doi:10.1016/j.exer.2005.03.006.

106. Escher, P.; Cottet, S.; Aono, S.; Oohira, A.; Schorderet, D.F. Differential neuroglycan C expression during retinal degeneration in Rpe65-/-mice. Molecular vision 2008, 14, 2126–2135.

107. Inatani, M.; Honjo, M.; Otori, Y.; Oohira, A.; Kido, N.; Tano, Y.; Honda, Y.; Tanihara, H. Inhibitory effects of neurocan and phosphacan on neurite outgrowth from retinal ganglion cells in culture. Investigative ophthalmology & visual science 2001, 42, 1930–1938.

