## Supplementary information for "Chromatin remodeling enzyme Snf2h is essential for retinal cell proliferation and photoreceptor maintenance"

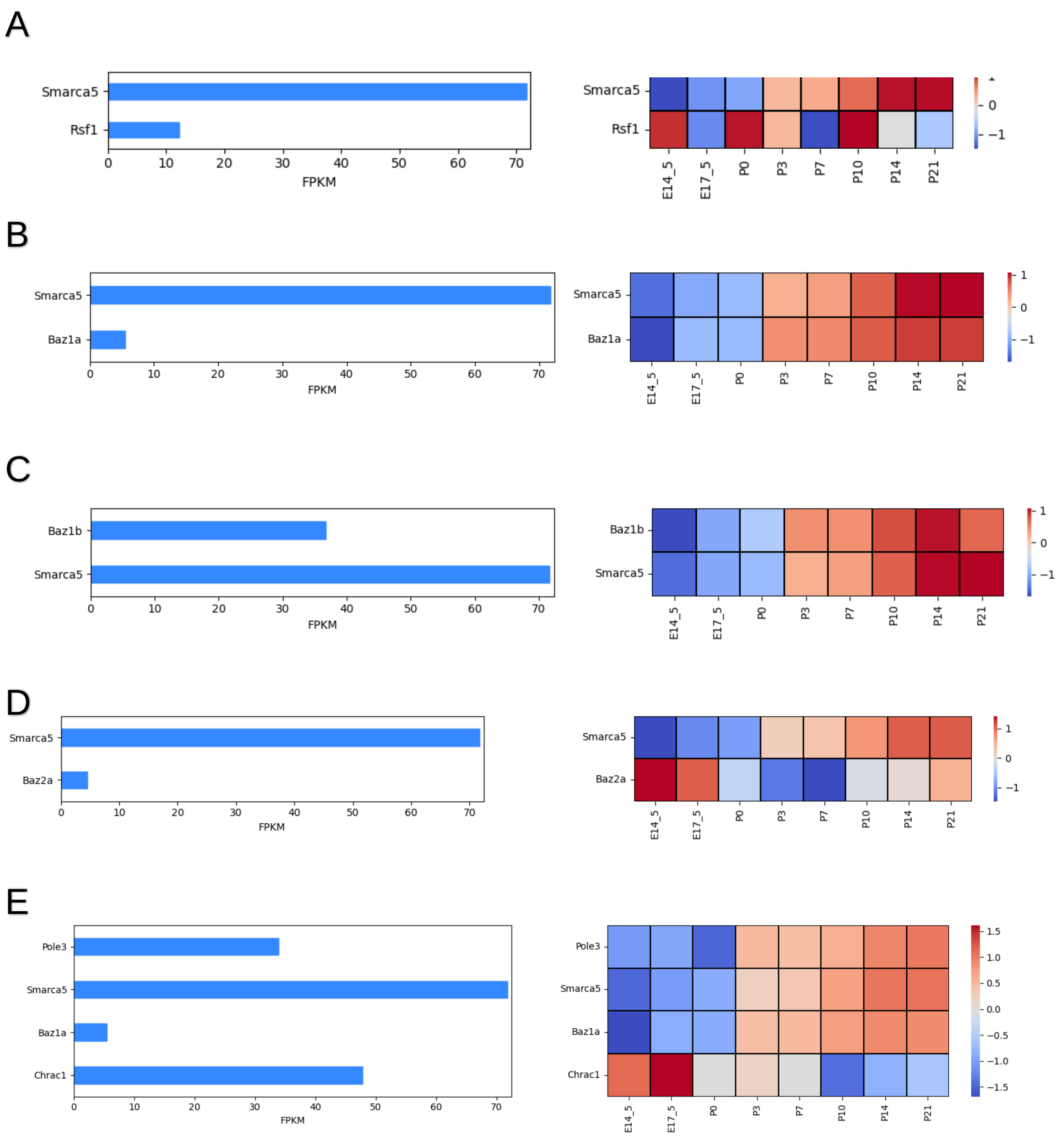


**Supplementary Fig. 1. Expression of components of ISWI complexes during retinal development.** Heatmap showing expression of components of RSF (A), ACF (B), WICH (C), NoRC (D), and CHRAC (E) complexes in retina at E14.5, E17.5, P0, P3, P7, P10, P14, and P21 (GSE87064 (Aldiri et al., 2017)).

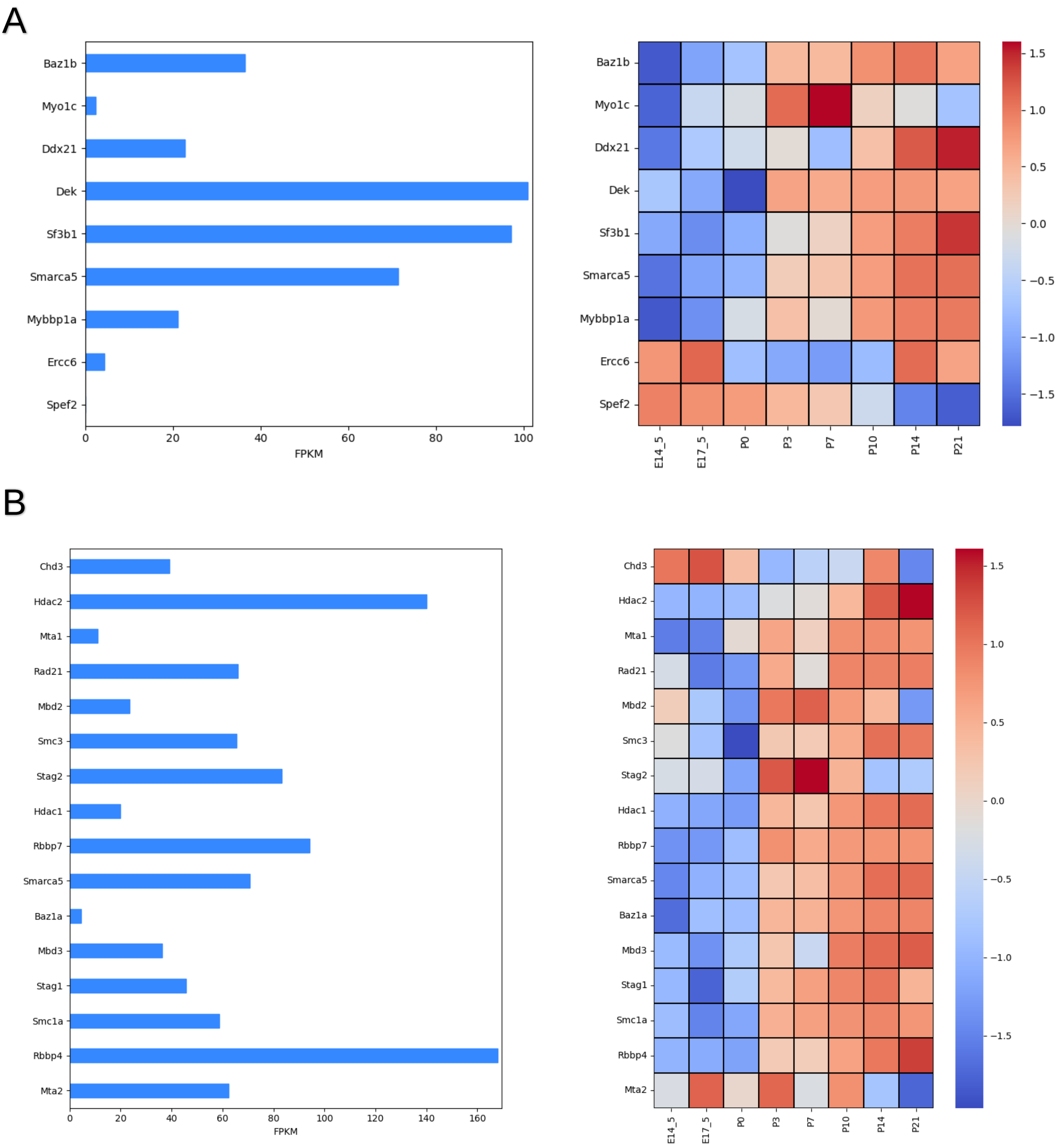


**Supplementary Fig. 2. Expression of components of ISWI complexes during retinal development.** Heatmap showing expression of components of B-WICH (A) and Cohesin (B) complexes in retina at E14.5, E17.5, P0, P3, P7, P10, P14, and P21 (GSE87064 (Aldiri et al., 2017)).

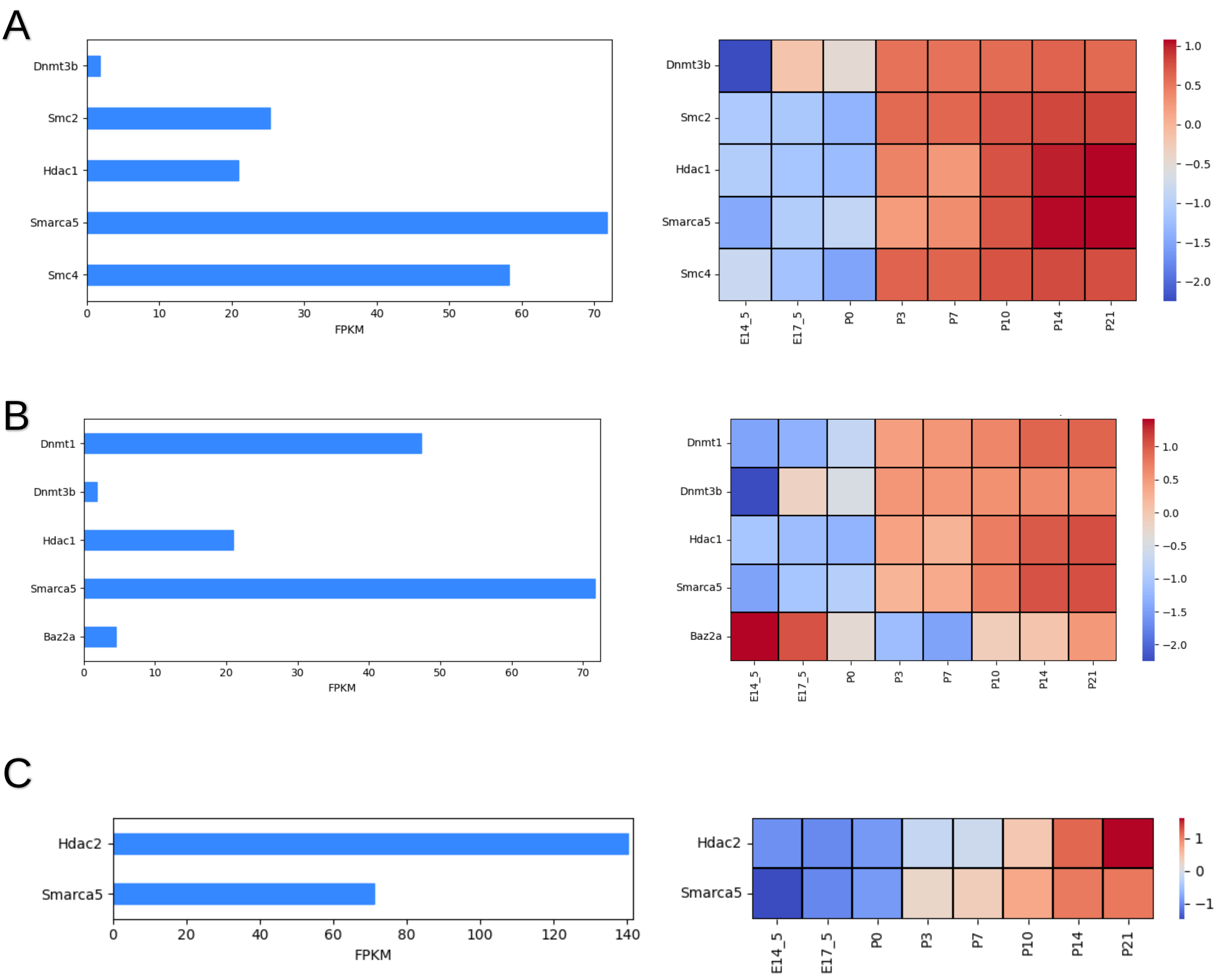


**Supplementary Fig. 3. Expression of components of Dnmt3b including complexes and Hdac2 during retinal development.**

Heatmap showing expression of components of Dnbt3b including complexes (A, B), and Hdac2 (C) in retina at E14.5, E17.5, P0, P3, P7, P10, P14, and P21 (GSE87064 (Aldiri et al., 2017)).

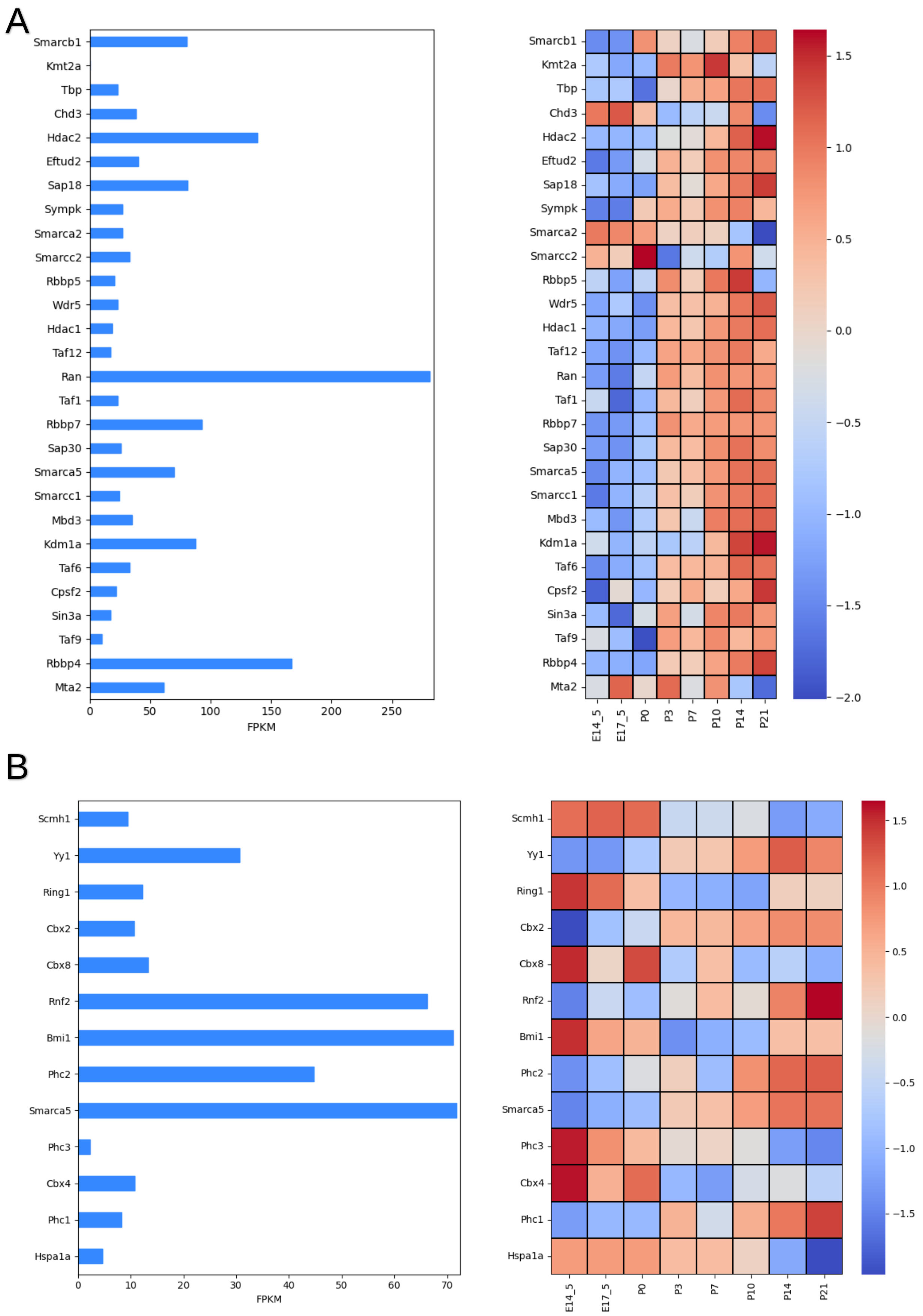


**Supplementary Fig. 4. Expression of components of All-1 and PRC1 complex during retinal development.**

Heatmap showing expression of components of All-1 (A) and PRC1 (B) complexes in retina at E14.5, E17.5, P0, P3, P7, P10, P14, and P21 (GSE87064 (Aldiri et al., 2017)).

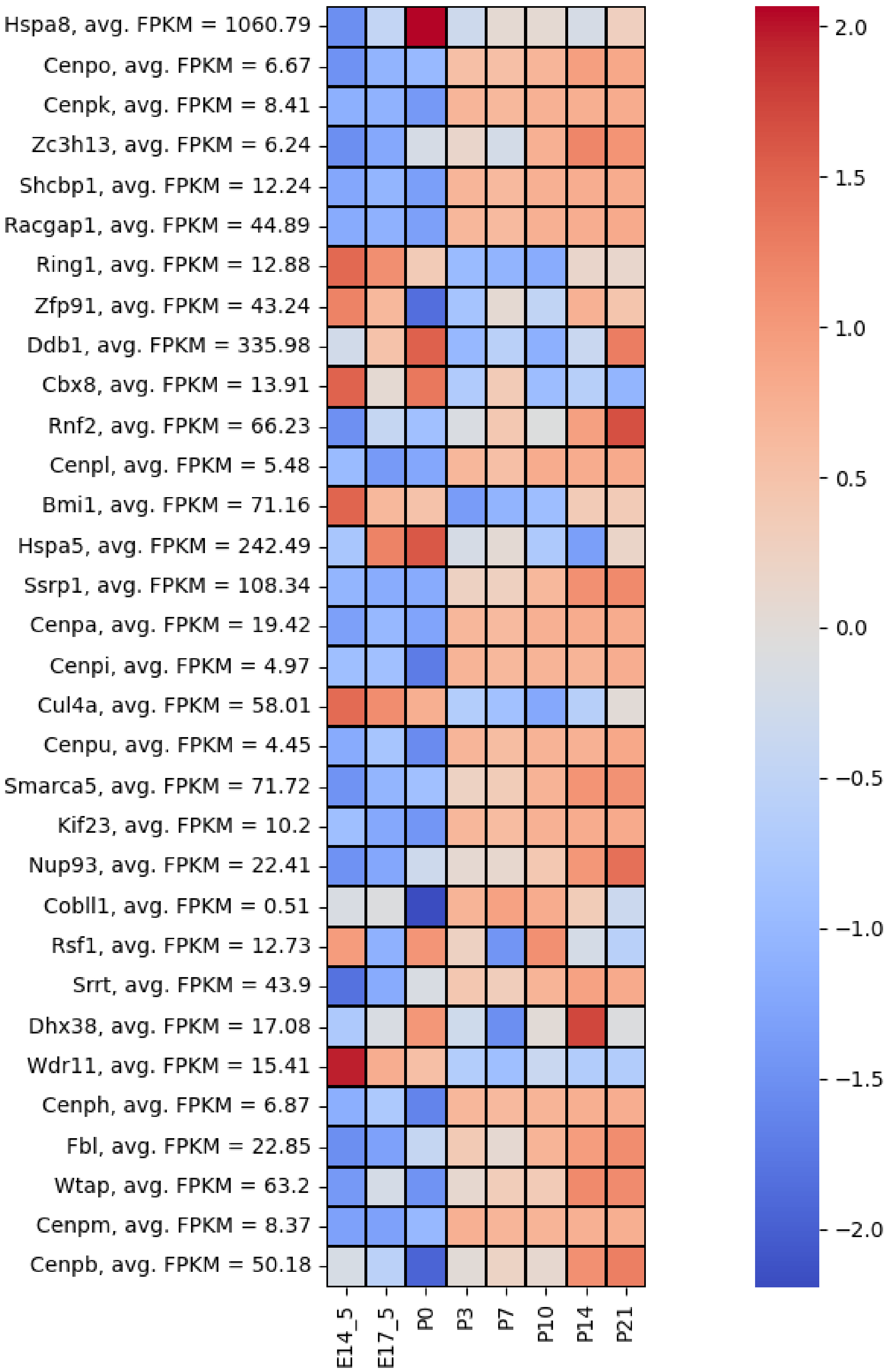


**Supplementary Fig. 5. Expression of components of centromere complex during retinal development.**

Heatmap showing expression of components of centromere complexe in retina at E14.5, E17.5, P0, P3, P7, P10, P14, and P21 (GSE87064 (Aldiri et al., 2017)).

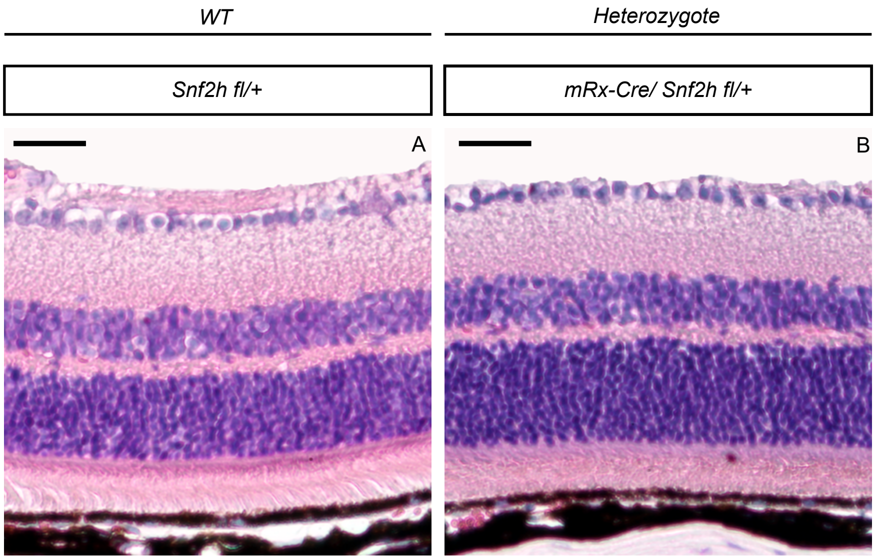


**Supplemental Fig. 6. Morphology of heterozygous mice at postnatal week 50 (PW50).**

Hematoxylin and eosin staining of wild-type (A) and mRx-Cre/ Snf2h$^{fl/+}$ (B) mice did not manifest obvious differences between the two genotypes. One active allele of the Snf2h gene is thus sufficient for the maintenance of an apparently normal gross retinal morphology.

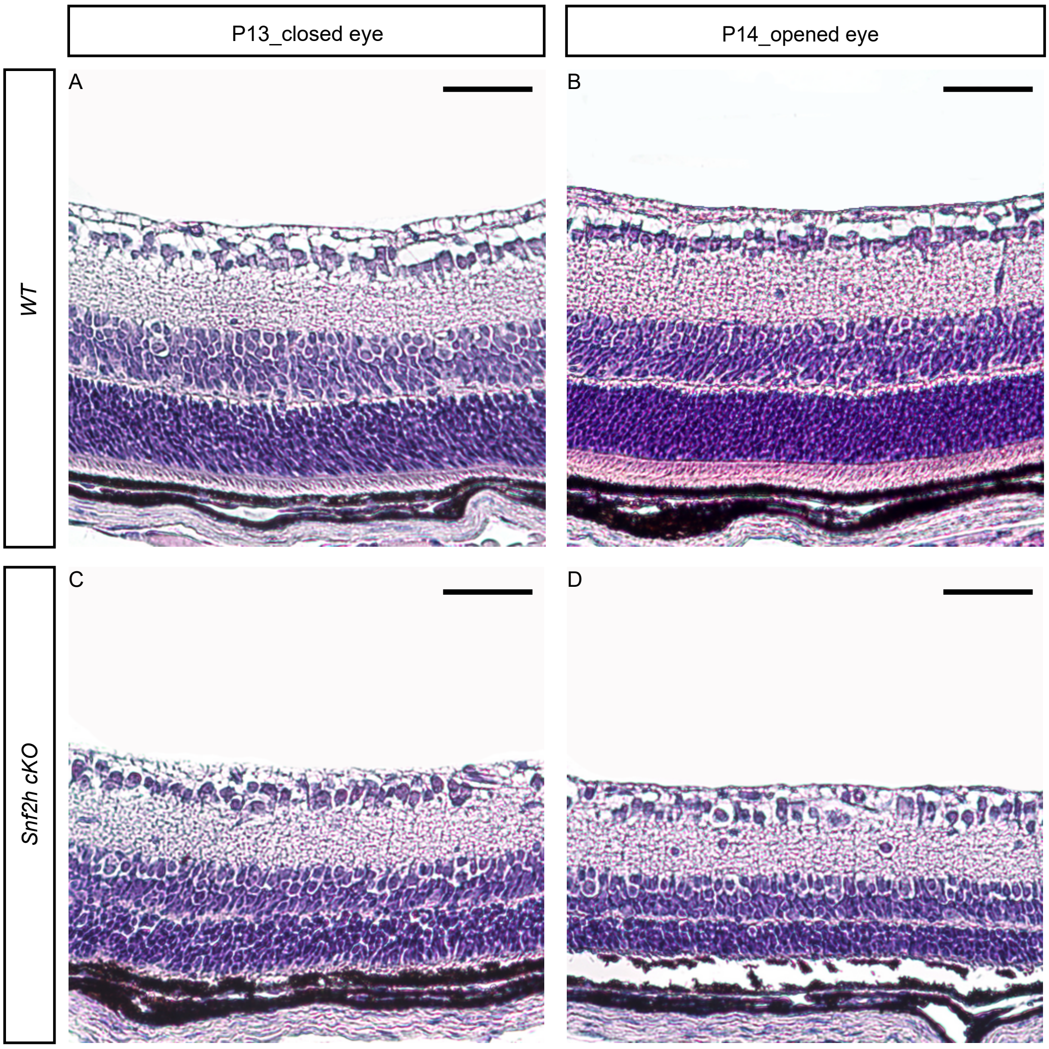


**Supplemental Fig. 7. Retinal morphology of Snf2h cKO at P13 and P14.**

Hematoxylin-eosin staining of postnatal retinal sections at postnatal day 13 (P13) and day 14 (P14) of wild-type and Snf2h cKO. The retinal section of Snf2h cKO at P14 (D) was clearly thinner compared with the same genotype at P13 (C).

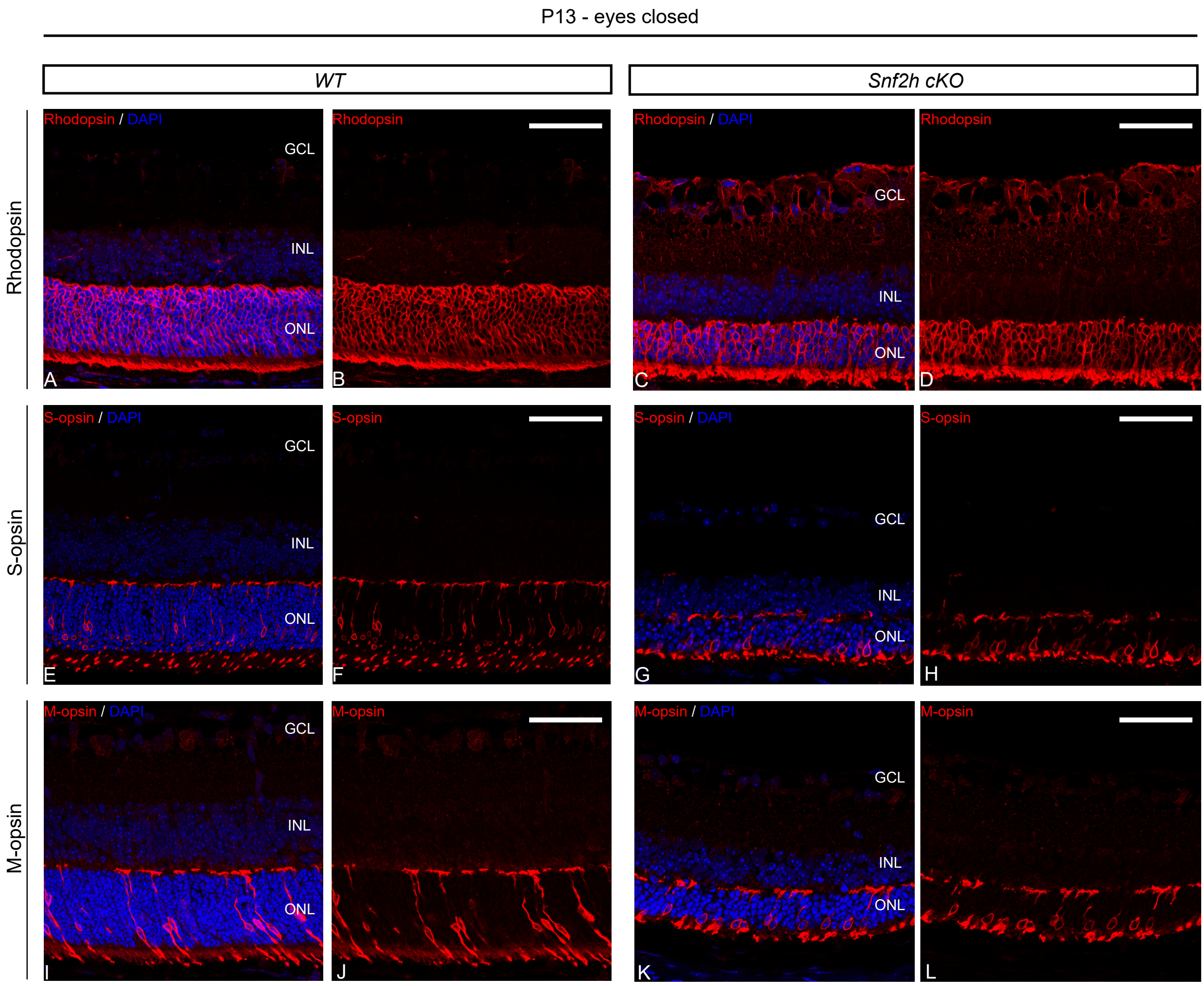


**Supplemental Fig. 8. Photoreceptor characterization at P13 (before eye opening).**

Rod photoreceptors in the wild-type retina (A, B). The rod photoreceptors in Snf2h cKO were present (C, D). Although the number of rods was reduced, their cell shape and positioning was comparable with the wild-type controls. The same result was obtained with cone photoreceptor staining (E-H, I-L).

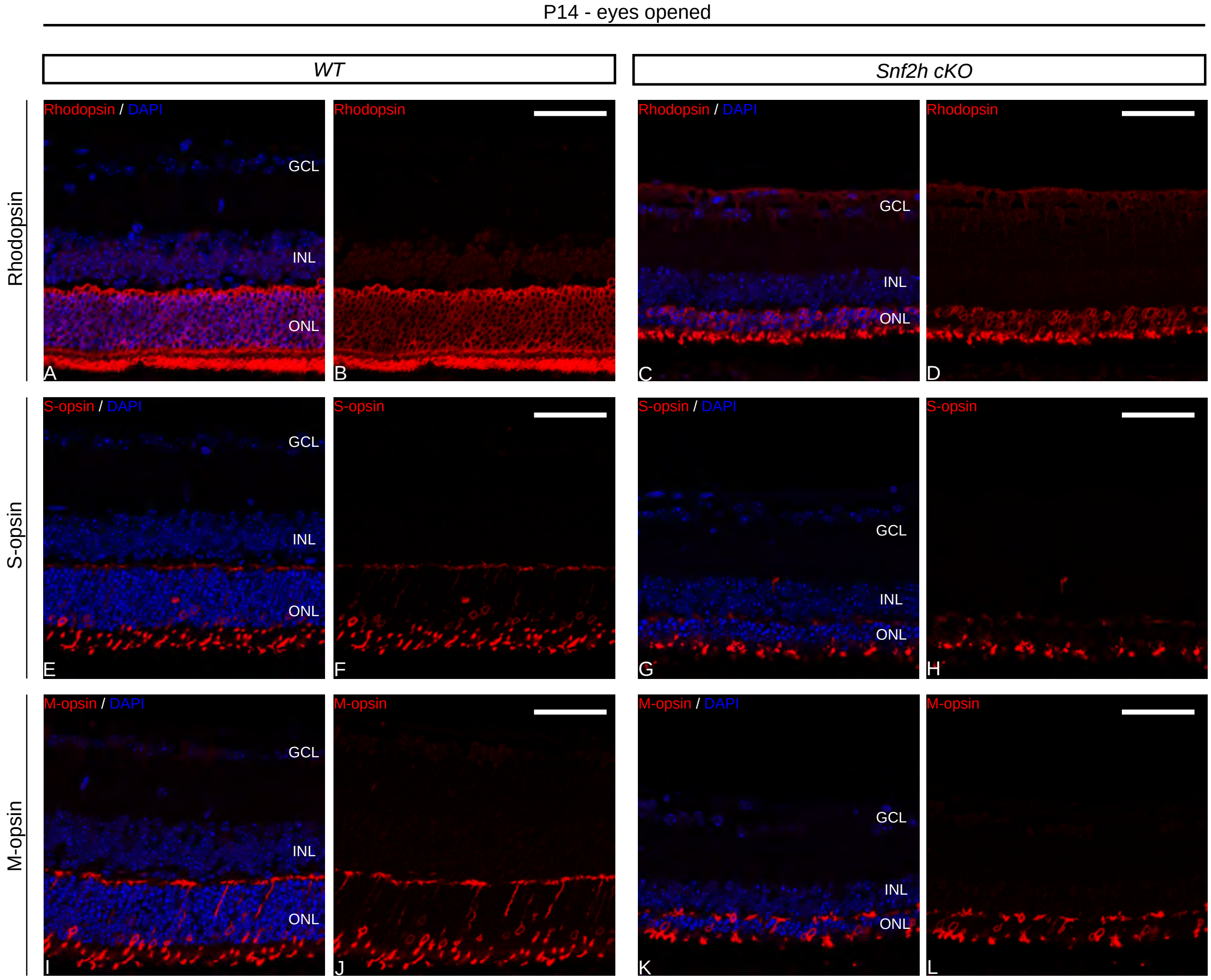


**Supplemental Fig. 9. Photoreceptor characterization at P14 (after eye opening).**

The rod photoreceptors in Snf2h cKO were preserved, but their length was reduced by about a half compared with control, which was associated with reduced thickness of the ONL in Snf2h cKO (A-D). The cone photoreceptors exhibited an abnormal shape, the outer segments were reduced, and the photoreceptor protrusions detected in wild-type animals inside the OPL were missing in Snf2h cKO (E-H, I-L).

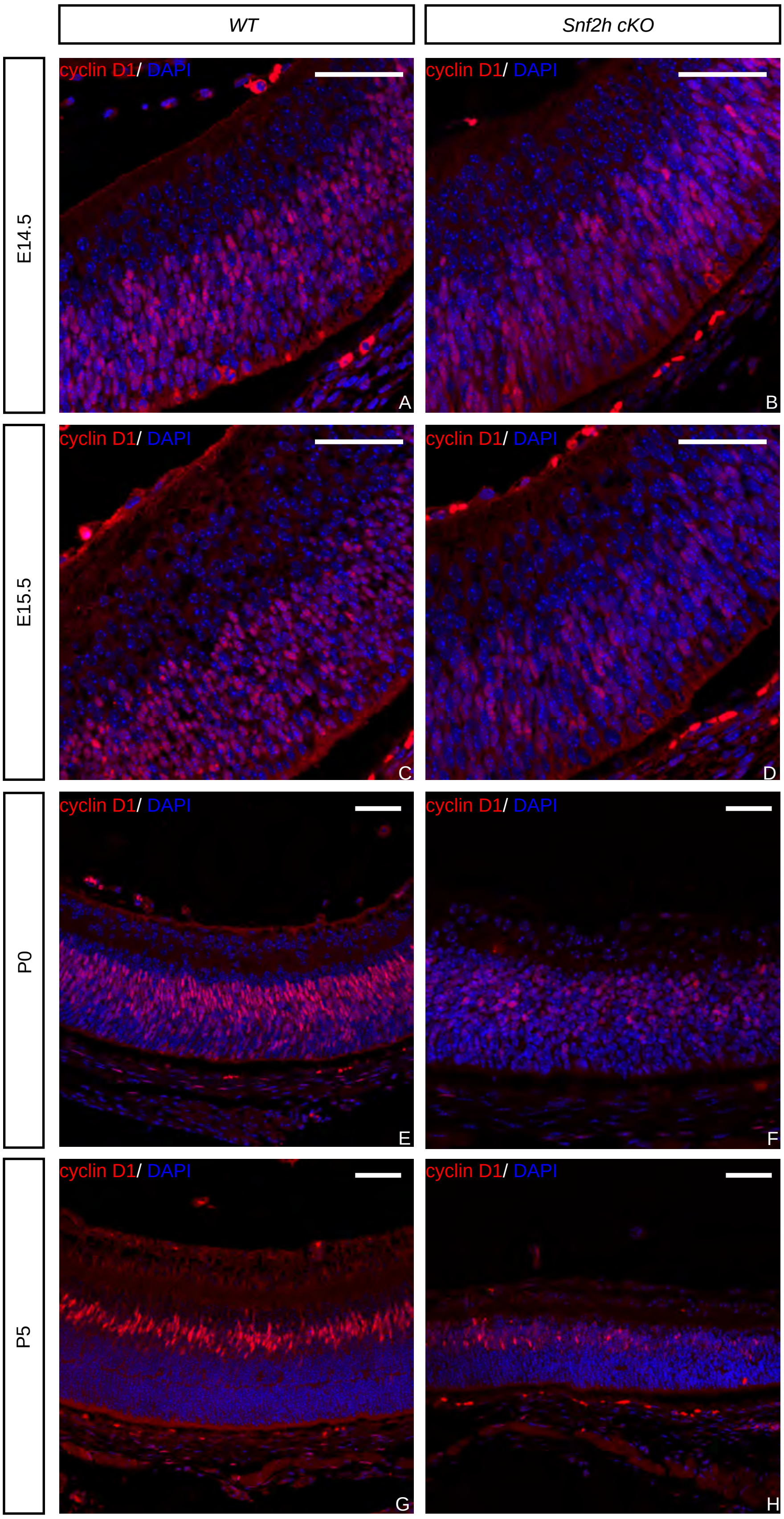


**Supplemental Fig. 10. Expression of cyclin D1 in Snf2h cKO during embryonic and postnatal stages.**

Immunostaining for cyclin D1, required for G1/S transition during the cell cycle, in wild-type and Snf2h cKO. Differences in cyclin D1 immunoreactivity appeared already during embryonic development (A-D). A dramatic decrease in cyclin D1-positive cells and cyclin D1 expression level in Snf2h cKO compared with wild-type controls was observed at birth (E, F) and at P5 (G, H).
